# Phosphate starvation stops bacteria digesting algal fucan that sequesters carbon

**DOI:** 10.1101/2024.04.07.588495

**Authors:** Yi Xu, Mikkel Schultz-Johansen, Huiying Yao, Isabella Wilkie, Leesa Jane Klau, Yuerong Chen, Luis H. Orellana, Finn Lillelund Aachmann, Mahum Farhan, Bowei Gu, Greta Reintjes, Silvia Vidal-Melgosa, Dairong Qiao, Yi Cao, Jan-Hendrik Hehemann

## Abstract

Algae provide a solar powered pathway to capture and sequester carbon by injecting stable fucan made from carbon dioxide into the ocean ^1–4^. Stability of the pathway is at odds with the presence of marine bacteria with genes of enzymes that can digest fucan and release the carbon dioxide ^5^. Biochemical explanations for stable fucan remain hypothetical ^6^. We assembled a biological carbon cycle model and found phosphate limitation enhanced fucan synthesis by algae, stopped digestion by bacteria and thereby stabilized the fucan carbon sequestration pathway. Marine microalgae *Glossomastix* sp. PLY432 increased synthesis of fucan, a part of its extracellular matrix, under nutrient-growth limiting conditions. Rate and extent of fucan digestion by a marine, isolated bacterium of the *Akkermansiaceae* family decreased with decreasing phosphate concentration. Phosphate starvation restricted bacterial growth rate, biomass yield and in turn increased the amount of stable fucan. Phosphate is universally required for growth but rare relative to glycan carbon in photosynthesis-derived ecosystems. The fact that phosphate is required for replication, transcription and translation explains why bacteria can digest gigatons of laminarin with a few enzymes, but not fucan during nutrient limited algal blooms. We conclude phosphate starvation constrains the ability of bacteria to digest fucan, which evolves to maintain stability around algal cells and consequentially also to keep carbon dioxide in the ocean.

## Main

Sulfated fucan was recently discovered to resist enzyme catalyzed hydrolysis and oxidation providing a pathway to contain carbon dioxide in the ocean ^2^. The Intergovernmental Panel for Climate Change, IPCC reproducibly analyzed and advised the climate crisis requires first the reduction of carbon dioxide emissions. Only second follows innovation to increase carbon dioxide removal and storage capacity ^7^. Diatoms ^2^ and brown algae synthesize an extracellular matrix ^8^ that contains fucoidan, a sulfated fucan with antibacterial ^9^, antiviral ^10^ and antifouling activity ^11,12^. Algal blooms accumulate dissolved and particulate fucan. The biological carbon pump transports particulate fucan within sinking particles into deeper waters and sediments burying the contained carbon dioxide for hundreds ^3^ to thousands of years^13^. Photosynthesis of fucan in the sunlit ocean, gravitational sinking, convection, transport into the mesopelagic, bathypelagic and sedimentation store carbon dioxide long term but only if the resistance of the fucoidan, fucan carbon sequestration pathway persists. Although slower than less complex glycans, enzymatic digestion of fucans has been detected in some oceanic regions including the Gulf of Mexico and sediments in some marine areas ^14–16^. Heterotrophic bacteria with fucanases are at relative low abundance relative to laminarin degraders detected in the ocean ^5^ questioning both, their ability to digest fucan effectively but also the long term stability of the fucan sequestration pathway.

To investigate what stabilizes and destabilizes the fucan sequestration pathway we assembled a biological carbon cycle model. The microalgae *Glossomastix* sp. PLY432, isolated from the Atlantic was obtained from the Roscoff culture collection and produced a viscous extracellular matrix composed of fucan. Once purified and structurally characterized, the fucan was used to isolate a marine *Verrucomicrobiaceae* bacterium from the North Sea capable to digest the fucan. Comparative genomics, physiology and environmental datasets show phosphate starvation promotes algae to synthesize fucan, stops bacterial digestion and thereby stabilizes the fucan sequestration pathway. The results are of global relevance because fertilization of photosynthetic organisms for enhanced biomass production and carbon dioxide removal can have the opposite effect when stimulating bacteria to digest and release the carbon dioxide contained within fucan and other complex glycans that sequester carbon.

### *Glossomastix* exudes fucan matrix

*Glossomastix* sp. PLY432 belongs to the Heterokontophyta including brown algae and diatoms that synthesize fucans. The *Glossomastix* genus contains one 18s rRNA sequence (GenBank: AF438325.1)^17^. For phylogenetic placement and validation we sequenced the V4-V5 region of the 18s rRNA gene. Phylogeny, supported by a maximum likelihood analysis, ordered the sequence into the Pinguiophyceae class within the Ochrophyta. The sequence was 96.31% pairwise nucleotide identical to *Glossomastix chrysoplasta* (**Fig. 1a**). Spherical *Glossomastix* cells of 4.9-8.5 µm size are encapsulated by an extracellular matrix **(Supplementary Table 1)**. Microscopy visualized a transparent, globular sphere with a radius of 3.4-6.4 µm housing one or two algal cells (**Fig. 1b and Extended Data Fig. 1a**).

**Fig. 1:**
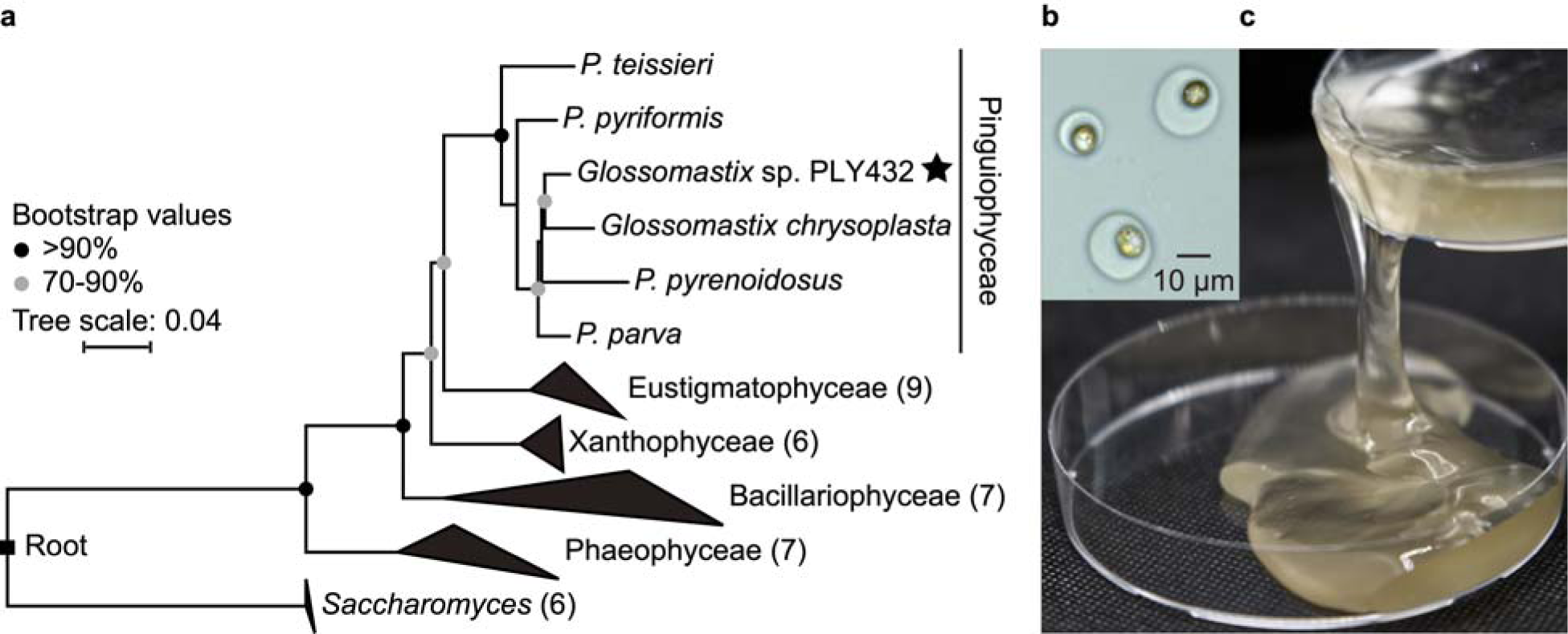
The heterokont microalga *Glossomastix* sp. PLY432 secretes a fucose-rich mucus that accumulates. **a,** Phylogenetic relationship of heterokont algae (class level) based on the V4-V5 region of 18S rRNA gene sequences. Numbers in parentheses indicate species contained in collapsed branches. *Saccharomyces* is added as an outgroup. Bootstrap values below 70% are not shown. **b,** Microscopy image with scale bar showing golden *Glossomastix* cells wrapped in transparent mucus. **c,** Decanting of a three month old *Glossomastix* culture reveals the viscous nature of fucose-rich mucus.

*Glossomastix* exude an extracellular matrix later identified as fucan. For almost a decade, we transferred algal cells to fresh medium every other week. Without dilution the solution became a weak gel, macroscopic viscosity and microscopic aggregates increased beyond three months (**Fig. 1c and Extended Data Fig. 1b**). This stable gel phenotype is conserved, in the North Sea ^2^, Adriatic Sea ^18^, around the globe ^19^ where fucose containing polysaccharides form microgels ^20^ that grow to become marine snow ^21–24^ or even stable macrogels ^25^ despite the presence of heterotrophic bacteria with genes of fucanases and fucosidases and other CAZymes ^5^. The culture included bacteria without consuming the visible abundance of fucan. Bacteria isolated from the culture did not digest the fucan. Instead gel strength increased with time indicating resistance to digestion. The algae produced a stable matrix without evidence of the fucan supporting bacterial growth.

### Nutrient limitation - fucan exudation

Not growing algae continue synthesis of fucan under nutrient limiting conditions. During two months of growth, the total glycan carbon content present in the secreted carbohydrates increased 36.42-fold, from 4.14 mg/L to 150.72 mg/L. Fucan synthesis increased after the growth phase, once the population reached carrying capacity when nutrients became growth limiting and cells stopped dividing (**Fig. 2a**). After they stopped dividing, we measured linear increasing concentrations of fucose, rhamnose and galactose, while galactose, glucose and glucuronate did not show this trend of linear increase (**Fig. 2b-c and Supplementary Fig. 1**). The constant number of cells and the linear monomer increase enabled calculating a net accumulation, potential synthesis rate for cells during the stationary phase. Microscopy of cells, viscosity of the solution and the abundance of glycan carbon indicate accumulation rate and secretion rate are similar. The carbon accumulation rate was 0.52 pg/cell/day, for fucose 14 µM/day (*r* = 1.00, *P* < 0.001), for rhamnose 4 µM (*r* = 1, *P* < 0.0001) and for galacturonate 1 µM (*r* = 1, *P* < 0.0001) (**Fig. 2c**), or ∼ 0.68 pg of fucose, ∼ 0.19 pg of rhamnose, and ∼ 0.06 pg of galacturonate per cell and day. Based on the volume and carbon content of diverse microalgae ^26–28^, we estimate a *Glossomastix* cell contains 24.49 pg carbon **(Supplementary Table 2)**. One cell exudes 2.12% of its carbon in form of carbohydrates, consistent with the fucoidan excretion rates of the brown macroalgae *Fucus vesiculosus* ^4^. The data shows a sustained population of algal cells sustains the fucan sequestration pathway.

**Fig. 2:**
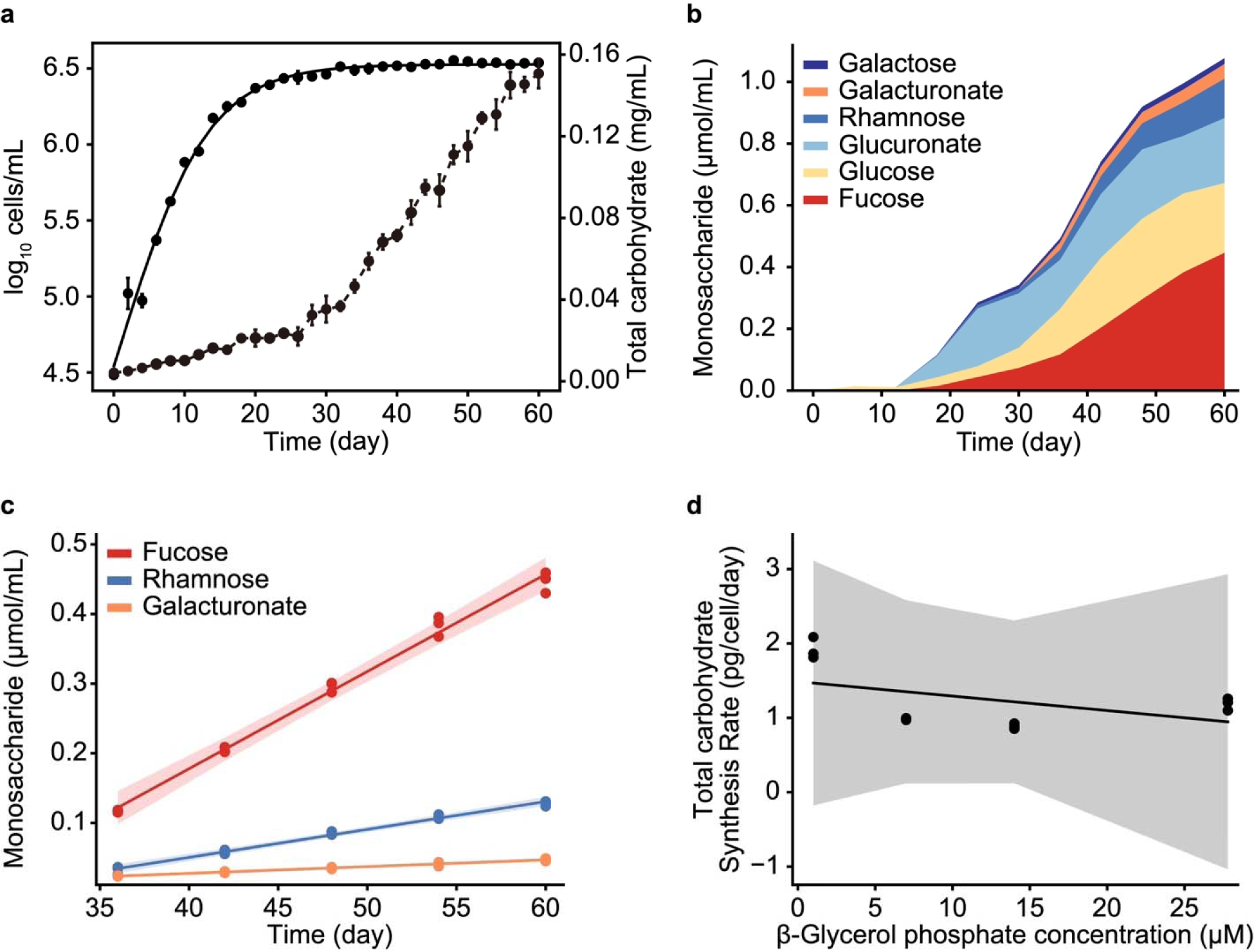
*Glossomastix* synthesizes and exudes fucan despite nutrient limitation. **a,** Monitoring of growth by cell counts (logistic fit, *R*^2^ = 0.9917) (solid line) and total carbohydrate content (dash-dotted line) in *Glossomastix* cultures. Experiments were performed as independent triplicates (n = 3) and error bars represent standard deviation of the mean. **b,** Monosaccharide composition of exudates collected from *Glossomastix* culture. Exudate samples were acid hydrolysed and monosaccharides quantified by HPAEC-PAD, each value was the mean of independent triplicate (n = 3). **c,** Linear regression analysis for monosaccharides in *Glossomastix* cultures (day 36 to 60, n = 3): fucose (slope = 0.014, *r* = 1, *P* < 0.001), rhamnose (slope = 0.004, *r* = 1, *P* < 0.0001) and galacturonate (slope = 0.001, *r* = 1, *P* < 0.0001). **d,** Correlation analysis of net carbohydrate synthesis rates of *Glossomastix* with different concentrations of phosphate (*r* = −0.48, *P* > 0.5). Data was generated from the stationary phase of *Glossomastix* culture with independent triplicate (n = 3).

Algae continued fucan exudation despite phosphate limitation. Experiments with diatoms and other microalgae found phosphate limitation promotes glycan exudation ^29,30^. We restricted access to phosphate and found the algae entered the stationary phase faster. Glycan synthesis increased during the stationary phase irrespective of the phosphate concentration (**Fig. 2a and Extended Data Fig. 2a-c**). Phosphate concentration neither affected the maximum, synthesis rate per cell and day (*r* = −0.42, *P* > 0.5) nor the growth rate (*r* = −0.89, *P* > 0.05) (**Fig. 2d and Extended Data Fig. 2d**). Continued synthesis and exudation of glycans sometimes referred to as overflow metabolism ^31–33^, has been verified with different algal species in laboratory and environmental settings and is consistent with the accumulation of fucan during algal blooms ^2,34^. A fucan carbon sequestration pathway that works with limited nutrients appears viable.

### Fucan structure

The structure of this fucan was required to verify its use as a model molecule. First, we purified the fucan from the *Glossomastix* exudate. Detailed fucan purification steps are outlined in Material and Methods and **Extended Data Fig. 3a**. A single, broad peak after anionic exchange and size exclusion chromatography in addition to one broad band on a gel proved the fucan was purified **(Extended Data Fig. 3b and Extended Data Fig. 4c**). We tested the purified fucan against four anti-fucan monoclonal antibodies (mAbs). These mAbs had been used in previous studies, showing they bind fucans from brown algae and diatoms and revealing the long term fucan carbon sequestration in marine sediments ^4,13^. Our enzyme-linked immunosorbent assay (ELISA) results showed mAbs BAM1 and BAM2 recognise the *Glossomastix* fucan (**Fig. 3a**). BAM2, binds a sulphated epitope in alpha configured fucan, while BAM1 binds a non-sulfated epitope ^35^. BAM2 showed higher binding signal to both our fucan and the positive control fucan compared to BAM1 (**Fig. 3a and Supplementary Fig. 2a**). However, this does not imply a higher abundance of BAM2 epitopes in our fucan molecule, because each antibody has a different affinity. BAM1 and BAM2 signals increased linearly with increasing fucan concentrations **(Supplementary Fig. 2b-c**).

**Fig. 3:**
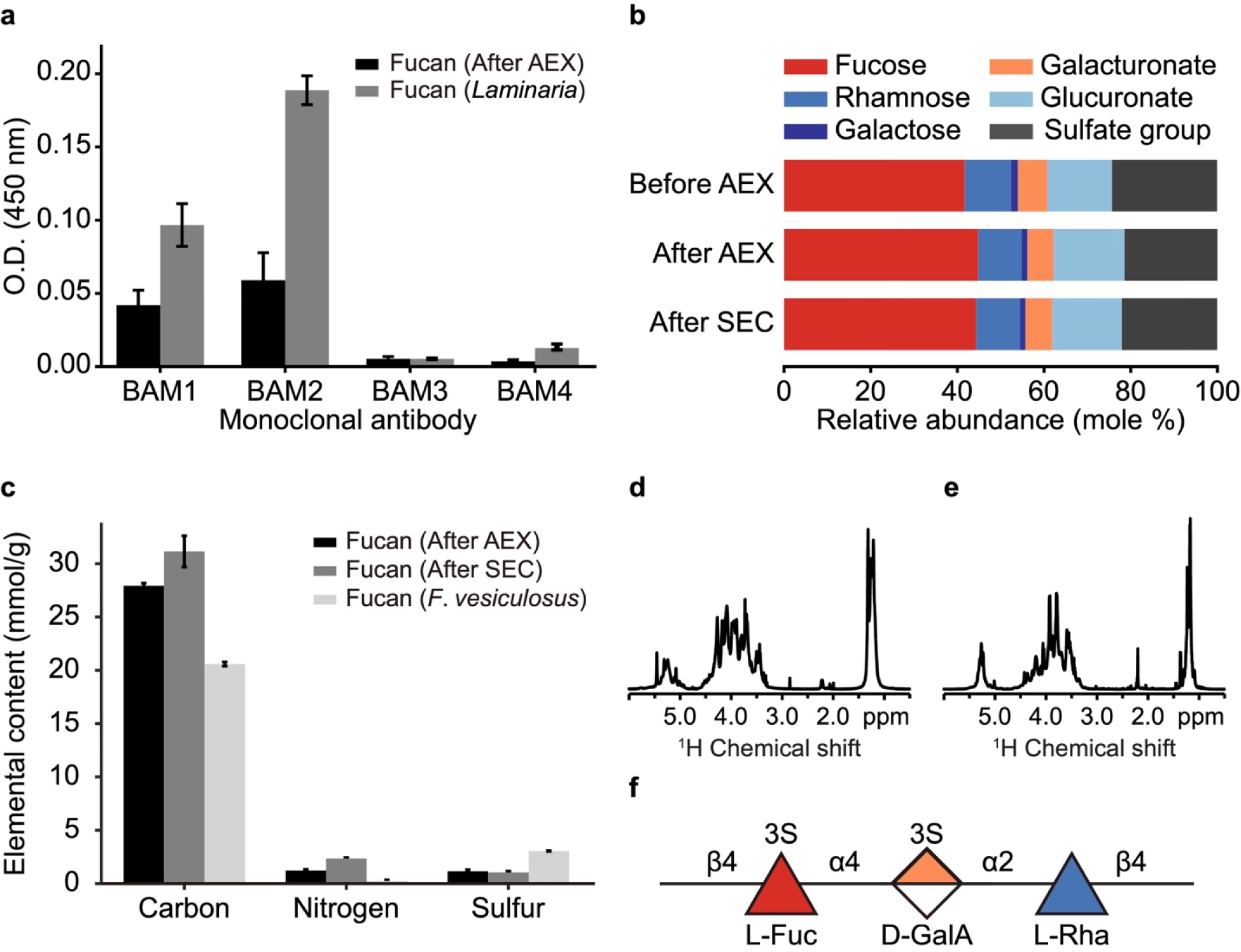
Antibodies, chromatography and NMR present a model structure for the fucan sequestration pathway. **a,** The binding of four fucan-specific monoclonal antibodies (BAM1-4) to AEX-purified *Glossomastix* fucan and to fucan from the macroalgae genus *Laminaria* (control) was assessed using ELISA. Fucans were used at 50 μg/mL on three technical replicates (n = 3). Optical density (O.D.). **b,** Monosaccharide composition of *Glossomastix* exudate (> 30 kDa) before and after purification with AEX and SEC. Glucosamine content less than 0.1% is not shown. Monosaccharide profiling was performed with HPAEC-PAD following acid hydrolysis of fucan and the data shown is the mean of triplicates (n = 3). **c,** Elemental analysis of purified *Glossomastix* fucan and fucan from *Fucus vesiculosus* (F8190, Sigma-Aldrich) on three technical replicates (n = 3), error bars represent standard deviation of the mean. **d,** 1D proton spectrum of purified fucan sample and **e,** 1D proton spectrum of desulfated fucan with water suppression. The samples were dissolved in D_2_O (200 µL, 99.96% D), spectrum recorded at 25°C and 800 MHz, ^1^H chemical shift internally referenced to the residual water signal (4.75 ppm). **f,** Structural representation of glycan fragment assigned by NMR spectroscopy of SEC purified *Glossomastix* fucan.

To know the building blocks of the fucan, we used compositional analysis by HPAEC-PAD. HPAEC-PAD analysis showed 20.41 ± 0.06 % (w/w) fucose, 3.33 ± 0.03 (w/w) galacturonate, 4.69 ± 0.02 % (w/w) rhamnose and 8.80 ± 0.06 (w/w) glucuronate. The molar ratio was 44.27% fucose, 10.18% rhamnose, 1.25% galactose, 6.11% galacturonate, 16.15% glucuronate and 21.98% sulfate (**Fig. 3b**). The composition was reproducible between batches from different years **(Extended Data Fig. 3c**). The elemental composition was 37.41% (w/w) carbon, 3.29% (w/w) sulfate and 3.33% (w/w) nitrogen. Monomer and elemental composition are comparable to fucans from diatoms and brown algae ^2,36,37^ (**Fig. 3c**). The reproducible composition enabled structure determination by nuclear magnetic resonance spectroscopy.

To add molecular resolution combinations of NMR spectra including HSQC, H2BC, IP-COSY, TOCSY and HMBC were used to resolve conserved parts of the structure. Desulfation reduced the complexity of the proton NMR spectra ^38,39^ (**Fig. 3d-e**). HSQC showed signals in the anomeric region A (δ_H_/δ_C_: 5.29; 103.0), B (δ_H_/δ_C_: 5.23; 103.0), C (δ_H_/δ_C_: 5.23; 100.7), and D (δ_H_/δ_C_: 5.01; 102.2) **(Supplementary Fig. 3a and Supplementary Table 3)**. Chemical shifts identified monomers and their configuration A: α-_D_-GlcA_p_; B: α-_L_-Fuc_p_, C: α-_D_-GalA_p_ and D: β-_L_-Rha_p_, inter-residue HMBC correlations, verified by NOESY correlations connected monomers with glycosidic bonds. The main chain is a repeating trimer of: −4)-α-_L_-Fuc*p*-(1-4)-α-_D_-GalA*p*-(1-2)-β-_L_-Rha*p*-(1-showing it is a heterofucan, hereafter the fucan (**Fig. 3f**). The α-_D_-GlcA*p* residue was not correlated in HMBC nor NOESY spectra, so it’s position and linkage remain unresolved.

The sulfated structure showed a higher chemical shift for proton (∼ 0.6 ppm) and carbon (∼ 7.4-8.0 ppm) from α-_L_-Fuc*p* (B’3) at position 3 (δ_H_/δ_C_: 4.52; 78.8) and α-_D_-GalA*p* (C’3) at position 3 (δ_H_/δ_C_: 4.69; 80.69) indicating sulfation on both of these monomers **(Supplementary Fig. 3b and Supplementary Table 3)**. Combined main signals indicate a trisaccharide of −4)-α-_L_-Fuc*p*(3-SO_3_)-(1-4)-α-_D_-GalA*p*(3-SO_3_)-(1-2)-β-_L_-Rha*p*-(1-(**Fig. 3f**). Acetyl CH_3_ groups were observed (δ_H_/δ_C_: 2.23; 23.3 and 2.21; 23.1) but not assigned to the trisaccharide. Minor signals indicate less abundant variations to the proposed trisaccharide core structure and potential branching. Variation is consistent with HPAEC-PAD finding relatively more fucose and rhamnose than galacturonate. Antibodies, chromatography and NMR revealed a structure related to other fucans but with a sequence of monomers we have not seen reported before. Previous work found a new structure for each fucan from another algal species ^40^. For example the fucoidans purified from six brown macroalge belonging to Fucales, Laminariales or Ectocarpales, all have alpha-configured fucose as main building block but differ regarding linkage connectivity, monomer composition, sulfation density and side chain modifications ^5^. Conclusively, this first, partial structure of a microalgal fucan is like all the others - different and therefore an equally perfect model molecule for the fucan sequestration pathway.

### Bacterium digests fucan

Bacteria with enzymes to digest fucan from microalgae remain unknown. Bacteria from the *Glossomastix* co-culture did not consume the fucan. Absence of bacteria with enzymes adapted to this fucan presents an alternative hypothesis for its stability. We tested bacteria in seawater and intertidal pore water from multiple sites along a beach on the island of Helgoland **(Extended Data Fig. 4a**). Bacteria from three sites consumed the fucan. After dilution into fresh medium, bacteria from two sites maintained the ability **(Extended Data Fig. 4b-c**). The results show bacteria evolved to digest it in the ocean and rejects the absence of adapted bacteria as a cause for stable fucan.

In a community, infection by other bacteria, phages, grazing, death by antibiotics, competition for resources, and many other unknown factors may render bacteria with adapted enzymes incapable to digest the fucan. Isolating a cause and effect requires an isolated bacterium. However, after obtaining isolated colonies from plates and reintroducing the clonal bacteria into liquid medium, they did not show the ability to digest the fucan. Only after reducing the concentration of fucan on the solid agarose plate did *Verrucomicrobiaceae bacterium* 227, hereafter *V*_227, begin to grow white, contrasting colonies **(Extended Data Fig. 4d-e**). *V_*227 was the only one digesting the fucan but only with presence of additional nutrients, trace elements and vitamins (**Fig. 4a**). These results indicate bacteria can digest the fucan but only when the environment provides enough additional resources and when the fucan is not too concentrated.

**Fig. 4:**
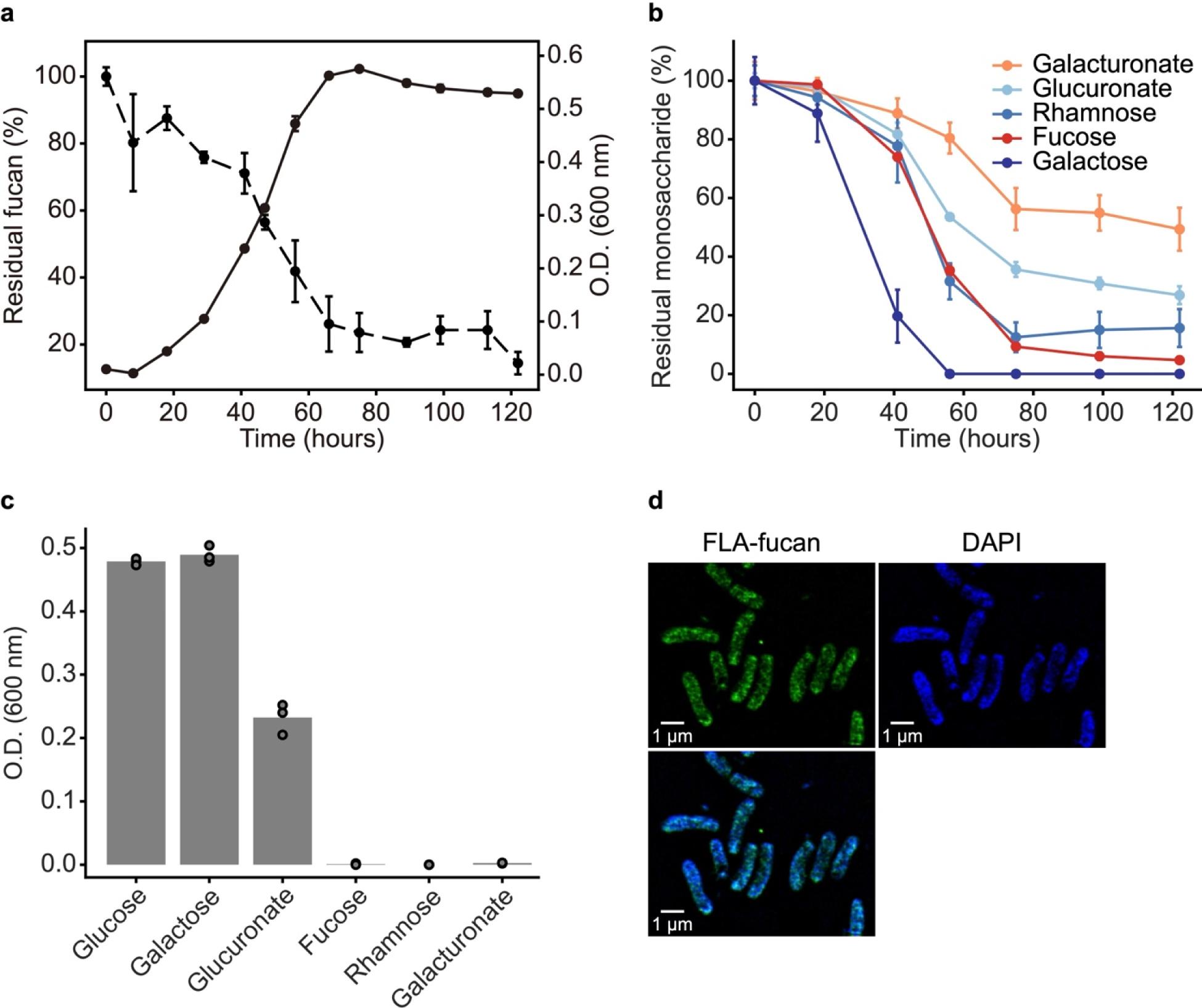
Selfish mode of fucan digestion indicates *Verrucomicrobiaceae bacterium* 227 (*V_227*) adapted to nutrient limitation. **a,** Growth of *V*_227 in MMT-YE medium with 0.05% (w/v) fucan (solid line) and the evolution of relative content of fucan in the culture supernatant (dash-dotted line). **b,** Evolution of concentration of total fucose, rhamnose, galacturonate, glucuronate and galactose in the culture supernatant. The experiment was performed in independent triplicate (n = 3), error bars are the standard deviation of the mean.**c,** Growth of *V_*227 with different monosaccharides as substrates. The experiment was performed in independent triplicate (n = 3). **d,** Selfish uptake of fucan by *V_*227. Super-resolution images of *V_*227 cells at day 3. DAPI (blue) signal shows nucleic acid. FLA-probe (green) signal was released from fluoresceinamine-labeled fucan. Merge DAPI signal and FLA-probe signal (Bottom of the panel).

Enough nutrients enabled *V*_227 to grow and digest 80% of the fucan indicating the two processes are coupled (**Fig. 4a**). Growth being coupled to digestion may seem obvious but becomes significant when considering algae can synthesize fucan without the nutrients required for growth ^4^. Another 5% was consumed by the bacterium during the stationary phase, after 122 hours 14.43% ± 3.35% of the fucan remained in solution. HPAEC-PAD analysis showed *V*_227 consumed 90.69% fucose, 87.51% rhamnose, 43.77% galacturonate, 100% galactose and 73.17% glucuronate but some of these monomers were consumed only when they were a part of the fucan polymer (**Fig. 4b**).

Selfish mode of glycan uptake suggests digestion of fucan may be influenced by nutrient limitation. *V_*227 did not grow with individual fucose, rhamnose, and galacturonate. Instead the fucan was imported and digested with enzymes inside the cell (**Fig. 4c**), an assumption consistent with the absence of oligosaccharides from the bacterial culture on a gel **(Extended Data Fig. 5**). We verified import of the fucan followed by internal digestion by fluorescently labeling the fucan to visually monitor the mode of consumption ^41^. Microscopy showed *V*_227 incubated with the FLA-fucan became fluorescent, and this is an established test for internal digestion, also known as selfish uptake (**Fig. 4d**). Selfish uptake is common for bacteria in intestines ^42^, rumen ^43^, ocean ^44^ and reduces loss of limited resources including secreted enzymes. Selfish uptake enables bacteria to conserve limited nutrients and therefore shows what they may be missing.

Next we asked whether the ability to digest this fucan is common. We compared *V*_227 with two other from the handful of known, characterized fucan-digesting bacteria. *V_*227 digests this fucan but not fucan from *Fucus vesiculosus* **(Extended Data Fig. 6a**). The *Flavobacterium Wenyingzhuangia fucanilytica* CZ1127^T^, digests fucoidan, fucan from sea cucumbers ^45^ but not this fucan. *Lentimonas* sp. CC4 digests six different fucoidans, fucans from six species of brown macroalgae with hundreds of enzymes, but not this fucan ^5^ **(Extended Data Fig. 5**). The specificity, exclusivity of these catalytic reactions verifies fucan structural diversity as a driver of enzyme diversity and confirms bacteria capable to digest fucans are rare.

### Genome phosphate limitation

Deoxyribonucleic acid, DNA sequencing of *V*_227 gave two contigs, one chromosome with 6,34 Mbp, and a megaplasmid with 0,42 Mbp. Phylogeny showed *V*_227 belongs to the *Akkermansiaceae* family ^46^. This family also includes the human commensal *Akkermansia muciniphila* that digests mucin, a fucan homolog, which is secreted by human, mammalian epithelial cells ^47^. Within the family *V_*227 belongs to the SW10 clade clades containing so far only sequences. *V_*227 shares 73.54% average nucleotide identity (ANI) with a metagenome assembled genome (WKH.23) from a river in Southern China ^48^ and is the first cultivated member strain within this genus (**Fig. 5**).

**Fig. 5:**
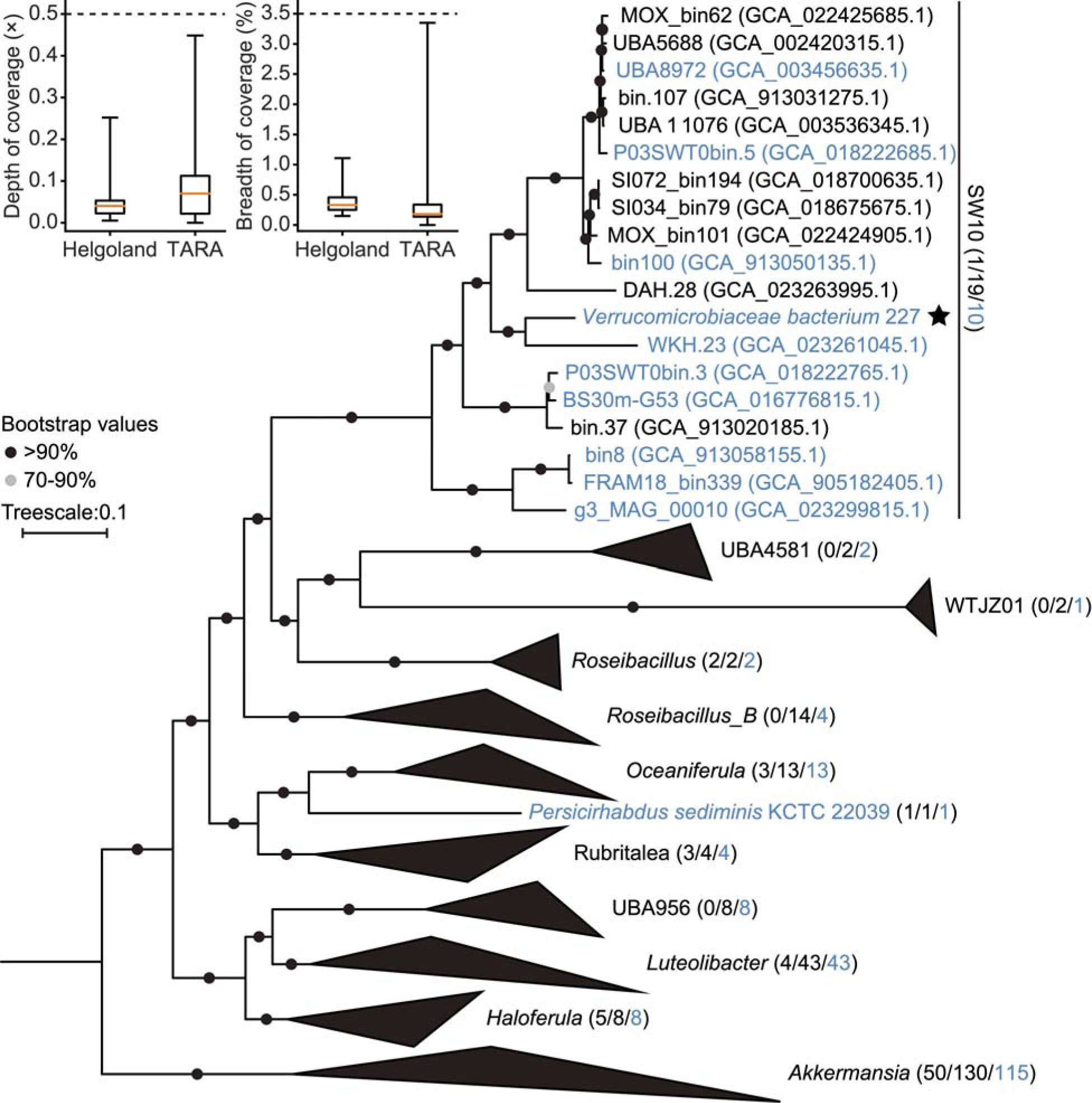
High affinity phosphate transporters common in glycan utilizing *Akkermansiaceae* reveals they adapt to environments with nM concentration of phosphate. The global abundance of *V_*227 in the ocean. Coverage depth and breadth in the TARA dataset and Helgoland spring bloom dataset. Below the dashed line indicates very low abundance in the dataset (Top of the panel). Phylogenetic tree based on 120 conserved genes from 246 selected *Akkermansiaceae* genomes (Bottom of the panel). Parentheses show number of culturable and number of total samples. The blue color indicates that the genome contains the high-affinity phosphate transport system (PstSABC), or the number of genomes containing this PstSABC system on each clade. *Verrucomicrobiaceae bacterium* 227 (*V_*227) is the only culturable bacterium in the genus SW10.

We used the genome sequence of *V_*227 to screen for its presence in global, circumnavigation TARA and Helgoland spring bloom datasets from the surface ocean. The depth and breadth of coverage were relatively low in the queried datasets e.g. < 0.5X and less than 3% (**Fig. 5**). Relatively low abundance of *Verrucomicrobiaceae* including their sulfatases, fucosidases and fucanases ^5^ remains intriguing considering fucose-containing glycans hold up to 20%, ∼ 20 µM of the dissolved organic carbon in the surface ocean ^49^. This low abundance may be related to phosphate concentration. Especially in oligotrophic regions of the surface ocean 0-300 nM, but also during later phases of coastal algal blooms phosphate rather than the dissolved organic carbon content restricts the growth of algae and bacteria ^50^. The phosphate concentration increases with depth until it reaches 1-3 µM in the bathypelagic ^51^. Recently the relative abundance of *Verrucomicrobiaceae* was found to increase in sinking particles that transport fucan into marine sediments ^52^. Hence, the phosphate concentration may influence the abundance, ability and distribution of fucan-digesting bacteria and therefore the stability of the fucan sequestration pathway.

Distribution and types of genes in the chromosome and plasmid indicate glycan digestion coupled to phosphate acquisition. The genome contains genes for the TCA cycle, and some genes for the ATP synthase are on the plasmid **(Supplementary Table 4-5)**. We identified about 100 carbohydrate active enzymes and sulfatases and found both enzymes on chromosome and plasmid **(Supplementary Table 4)**. The annotations are consistent with glycans digested by *V*_227 **(Extended Data Fig. 6b-c**). Transporters for acquisition of phosphate and other nutrients are located on chromosome and plasmid. The genome contains 206 genes annotated as membrane transporters, including 8 TonB-dependent receptors. TonB-dependent transport proteins require energy to import ferric chelates, vitamin B_12_, nickel complexes, peptides and oligosaccharides ^53^. 12 major facilitator superfamily transporters import oligosaccharides, monosaccharides and amino acids ^54^. The chromosome also contained 3 Na^+^/phosphate symporters (NptA) with low affinity for nutrient-rich environments ^55,56^. For phosphate-deprived environments there are two homologs of high affinity ABC-type phosphate transport system composed of four proteins in an operon (PstS,A,B,C). One operon is located on the chromosome and the other on the plasmid. One regulatory protein PhoU, for regulation of the PstSABC proteins, may regulate the two operons **(Supplementary Table 6)**. Notably, homologs of this high affinity system are present in 211 of the 246 genomes shown in the tree (**Fig. 5 and Supplementary Table 7)**, including the intestinal *Akkermansia muciniphila* suggesting phosphate starvation may be common not only in the ocean.

The PstSABC system has an affinity constant between 200-400 nM to bind, import phosphate at nM concentration suggesting phosphorus limits the digestion of complex glycans during algal blooms ^56^. Accordingly, during North Sea algae bloom the ABC transporter, proteins expression for phosphate and phosphonate uptake increased due to competing algae and bacteria that depleted the phosphate pool ^57^ while fucan remained stable and therefore accumulated ^2,34^. The nM phosphate uptake system also occurs in intestine and soil **(Supplementary Table 8)** including *Bacteroides thetaiotaomicron* ^58^, C*lostridium perfringens* ^59^ or other bacteria digesting the mucus matrix around human epithelial cells ^58^. The prevalence of phosphate transporters with nM affinity indicates that glycan-digesting bacteria from different environments experience phosphate starvation. Conclusively, phosphate starvation may constrain the ability of bacteria digesting complex glycans that surround and protect ^60^ eukaryotic cells.

### No phosphate no digestion

To test whether phosphate has an effect we increasingly reduced the concentration and monitored growth rate with laminarin or *Glossomastix* fucan as sole carbon source. Due to its biogeochemical and ecological relevance ^61^ we used the less complex laminarin as a control glycan. For laminarin, the growth rate remained relatively constant at 0.15 h^−1^ between low 1 µM and 50 µM phosphate concentration (*r* = - 0.76, *P* < 0.05). With fucan, the growth rate was overall lower and decreased linearly from 0.13 h^−1^ at 50 µM to 0.06 h^−1^ at 1 µM to phosphate concentration (*r* = 0.98, *P* < 0.001) (**Fig. 6a**). At 50 µM phosphate concentration, both glycans enabled similar growth rates, we stopped increasing the concentration as the growth rates plateaued. The convergence of growth rates shows the effect is not detectable in bacterial growth media that have higher phosphate concentration. The lowest phosphate concentration of 1 µM tested here is still higher than that in nutrient limited algal blooms ^62,63^. Yet, at 1 µM phosphate the mean growth rate was 2.98 fold slower with fucan than with laminarin. Intrigued by this observation we tested two other glycans, pectin and xylan **(Extended Data Fig. 7**). Similar to laminarin the growth rate did not decrease together with the decreasing phosphate concentration. The data shows for unknown reasons only fucan slows the bacteria significantly down compared to laminarin and other glycans when the cells are phosphate starved.

**Fig. 6:**
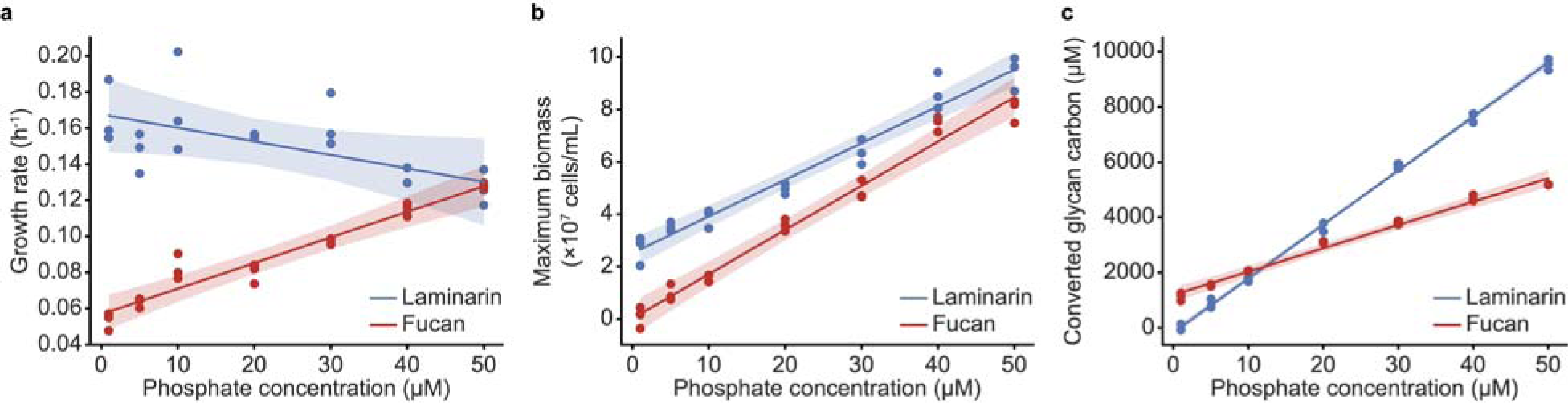
Phosphate starvation stops bacterial digestion enabling fucan carbon sequestration. **a,** Correlation analysis between growth rate and phosphate concentration on laminarin (*r* = −0.76, *P* < 0.05) and fucan (*r* = 0.98, *P* < 0.001). The fitting was performed using the mean growth rate. **b,** Correlation analysis between maximum biomass and phosphate concentration on laminarin (*r* = 0.99, *P* < 0.0001) and fucan (*r* = 0.99, *P* < 0.0001). The fitting was performed using the mean of maximum biomass. **c,** Correlation analysis between converted glycan carbon and phosphate concentration on laminarin (slope = 195.46, *r* = 1, *P* < 0.0001) and fucan (slope = 84.43, *r* = 0.99, *P* < 0.0001). The fitting was performed using the mean of consumption of polysaccharides carbon. The experiment was performed in independent triplicate (n = 3).

Next we investigated the relationship between biomass yield and phosphate concentration. Growth yield curves showed the biomass decreased with the phosphate concentration for laminarin and fucan. However, at phosphate concentration of 10 µM and lower the fucan significantly decreased the biomass compared to laminarin **(Fig.6b and Extended Data Fig. 8-9**). With fucan at 1 µM phosphate we count (1 ± 4) × 10^6^ cells/ml (linear model: 1.88 × 10^6^) and with laminarin (27 ± 6) × 10^6^ (linear model: 26.54 × 10^6^), a 27 fold difference (linear model: 14.13). Using the 14.13 fold difference from the linear model and assuming the phosphate was quantitatively converted into bacterial biomass, a cell growing with fucan contained 0.53 fmol and a cell with laminarin 0.04 fmol of phosphate **(Supplementary Table 9)**. These numbers fall within the range measured for different phyla of heterotrophic bacteria from different environments and in different growth phases ^64^. Using the conservative calculation fucan requires roughly ten times more phosphate to support the same number of cells. The converse conclusion is cells grown with laminarin can allocate ten times more phosphate to replication and other biochemical reactions that require phosphate. These physiological data provide a direct mechanism explaining the stability and accumulation of fucan during algal blooms. Laminarin stimulates faster growing bacteria (*Gammaproteobacteria*, SAR11, flavobacterial *Polaribacter* spp. and alphaproteobacterial *Rhodobacterales* spp.) ^57^ that rapidly deplete the phosphate and phosphonate pools, leaving not enough for Verrucomicrobiota and thereby limiting their abundance by restricting their ability to grow and digest fucan ^2,61^. The question remained does phosphate starvation stabilize fucan and other complex glycans?

Quantitative glycan carbon accounting finds fucan more stable with limited phosphate. To measure the conversion of glycan carbon relative to phosphate we quantified the glycan parts that remained in a given phosphate concentration. Residual fucan and laminarin carbon was quantified by HPAEC-PAD analysis of their constituent monosaccharides. The initial glycan carbon concentration minus the residual glycan carbon was used to calculate how much glycan carbon was converted per phosphate by the bacterium. The data show a fixed ratio of glycan carbon converted per phosphate. For 1 µM phosphate, 195.46 µM laminarin carbon (*r* = 1, *P* < 0.0001) (**Fig. 6c**) was converted. In case of the fucan, for 1 µM phosphate the bacterium converted significantly less carbon, 84.43 µM (*r* = 1, *P* < 0.0001). The data indicates for both glycans, carbon above the linear ratio, line remained stable and carbon below was converted. To verify we tested pectin and xylan and found growth yield and carbon conversion was for all four tested glycans constrained by the amount of phosphate **(Extended Data Fig. 7**). Comparing growth yield (**Fig. 6b**) with carbon conversion (**Fig. 6c**) shows the bacterium converts significantly more laminarin than fucan carbon to yield comparable bacterial biomass. Theoretically, glycolysis of glucose and galactose generates, before the TCA cycle, 2 ATP compared to 1 ATP from fucose, rhamnose, glucuronate, and galacturonate. Glucuronate and galacturonate yield 0 NADH where the other monomers yield two **(Extended Data Fig. 10**). Less biomass for more laminarin and carbon energy indicates loss. The TCA cycle rapidly extracts energy and releases carbon dioxide from glucose. Even without knowing what we lost the fucan is twice as stable and therefore of superior quality to store carbon.

Combined results show glycan types have different qualities for carbon sequestration and also provide ways to quantify this quality. The linear relationship of converted glycan carbon per phosphate (**Fig. 6c**) indicates a structure specific stability index. The index may change for example in a dynamic microbiome, where diverse bacteria compete for the same inorganic nutrients ^65^ that consequently fluctuate and sometimes restrict or stimulate growth and by extension the digestion of complex glycans ^66^. Glycan structural complexity or stability evolves on the surface of algae and other eukaryotes in the ocean, soils, intestines, around cells of plants. In all of these environments diverse types of cells including prokaryotes compete for the limited amount of phosphate ^67^, which is required for growth and to digest complex glycans. The index might be useful to quantify the stability of glycans in the presence of microbes for carbon sequestration, as prebiotics, as materials and as emerging drugs.

## Conclusion

Aligned growth rates between eukaryotes and prokaryotes are a fundamental tenet for sustainable co-existence of species, a balance easily disrupted by resource imbalance. Bacteria have smaller cells with more surface area for receptors and transporters to obtain nutrients from the environment ^68^. Algae compete with heterotrophic bacteria for nutrients and are except for carbon dioxide at a disadvantage. Support for this hypothesis comes from nutrient fertilization experiments conducted with ships and mesocosms across the ocean. Ocean fertilization with nutrients can stimulate algal blooms ^34,69^. However, experiments in the Arctic ^70^, Mediterranean ^50^ and Southern Atlantic Ocean ^71^ found addition of glucose, phosphate or iron stimulated heterotrophic bacteria. The scientists who conducted the experiment in the Arctic proposed the addition of glucose enabled heterotrophic bacteria to consume limited nutrients faster than algae shifting the system away from photosynthesis and carbon capture towards heterotrophy and release of carbon dioxide. Ocean and laboratory experiments indicate the carbon/nutrient ratio but also the complexity of the molecule that provides carbon energy can tune the balance between heterotrophic bacteria and algae. So how do algae persist? Pioneers of quantitative glycan accounting found algae derived, fucose containing glycans of unknown structure are stable for years in the global surface ocean^49^. These glycans hold 20%, ∼ 20 µM of the dissolved organic carbon in this for the largest part phosphate deprived (0-300 nM) system. Previously we thought complex glycans increase the growth of heterotrophic bacteria from the ocean to the human gut ^72^. Here we present fucan that significantly reduced bacterial growth rate and yield of bacteria potentially giving algae more time to access limited nutrients. When heterotrophic bacteria consume structurally simpler carbon energy compounds including laminarin and metabolites ^73^ they rapidly grow by consuming the essential nutrients. These nutrients are now missing for slower bacteria decreasing their ability to digest the fucan. Limited nutrients stabilize the fucan matrix around the algal cells physically excluding and biochemically constraining bacteria. Consequentially, when the fucan matrix is the primary carbon, it increases the space and time available for algae to access limited nutrients. We conclude these algae likely persist because of fucan that sequesters carbon in the ocean.

### Limitations

We studied here a limited carbon cycle consisting in essence of one alga, one bacterium and one glycan. Estimates predict 12000 to 30000 species of diatoms ^74–76^ are not alone with all the other algal phyla in the ocean. Glycan analysis finds a new fucan structure for each additional, studied algal species. Each change to a glycan structure requires a bacterium to adapt by acquisition or modification of a unique enzymatic cascade. We acknowledge one algal species, one bacterium and one fucan are a limited carbon cycle model. However, the here described organisms, molecules, results are consistent not only with ocean measurements but also with emerging basin scale modeling data of of microbial abundance, activity and inactivity ^77^. Measured and modeled data show compared with other taxa the growth of *Verrucomicrobiaceae* is relatively restricted in the surface ocean, in line with phosphate starvation as a cause for fucan stability. Conclusively, we have no knowledge that makes us think the here presented organisms, molecules and results are exceptional.

### Sustainability and inclusion statement

The here conducted experiments, methods, instruments, organisms are broadly accessible and therefore this work is inclusive for many scientists around the globe. The most advanced instrument was a nuclear magnetic resonance machine. The next advanced instrument was a HPAEC-PAD machine for detection of glycans with pulsed amperometric detection. HPLC machines with other modes of detection can also work for the analysis of glycans. Other commonly accessible instruments include a spectrophotometer. Only one bacterial genome was sequenced for this study limiting the amount of data that requires long term storage and sustainability. The limited amount of data we produced are available as excel sheets provided together with this manuscript.

## Materials and Methods

### Cultivation of *Glossomastix*

The microalgae strain, *Glossomastix* sp. PLY432, was obtained from the Roscoff Culture Collection (RCC3688) and cultivated in NEPCC medium (MediaDive: 1724) ^78^ at 15°C, with 140 µmol m^−2^ s^−1^ photosynthetic photon flux density provided by cool-white fluorescent lamps under a 12 h/12 h light/dark cycle. Pre-cultures grown for two weeks were used to inoculate fresh NEPCC medium. For large-scale cultures, 1 L culture was grown in 2 L Fernbach flasks with 20 mL pre-culture. For small-scale cultures, 150 mL NEPCC medium in 250 mL cell culture flasks was inoculated with pre-culture to an initial concentration of 3.05 × 10^4^ cells mL^−1^. Growth and carbohydrate production of *Glossomastix* was monitored over 60 days by sampling of 2 mL mixed culture every second day, or every forth days when *Glossomastix* was grown in phosphate-limited medium. Phosphate-limited NEPCC medium was obtained by adjusting the final concentration of β-glycerol phosphate. The initial concentration of algal cells in phosphate-limited cultures was 2.25 × 10^4^ cells mL^−1^. Cell counting was performed with a Neubauer haemocytometer (Marienfeld-superior, Cat No. 0640111) using 10 µL culture. Total carbohydrate was determined using 200 µL culture as described below. The remainder of the 2 mL samples were stored at −20°C for total monosaccharide determination. After three months of growth, *Glossomastix* cultures were sampled from the top and bottom part of the culture flask to assess low and high viscosity fractions, respectively. Cell morphology was examined with an EVOS FL Auto Microscope (Thermo fisher) and the size of mucus layers was evaluated manually using ImageJ ^79^.

### Phylogenetic analysis of *Glossomastix*

*Glossomastix* sp. PLY432 genomic DNA was extracted using the DNeasy Blood and Tissue Kit (Qiagen). The V4-V5 region of the 18S rRNA gene was amplified from purified genomic DNA using the 18S universal primers (574F-CGGTAAYTCCAGCTCYAV and 1192R-CAGGTGAGTTTTCCCGTGTT) in Q5^®^ High-Fidelity 2 × Master Mix according to the manufacturer’s instructions ^80,81^. PCR products were then purified with the QIAquick^®^ PCR Purification Kit (Qiagen) and sequenced at Eurofins Genomics (Cologne, Germany). The 18S rRNA genes of 34 species from 5 classes of the Ochrophyta (Pinguiophyceae, Eustigmatophyceae, Bacillariophyceae (Diatom), Phaeophyceae (Brown algae) and Xanthophyceae) were obtained from NCBI and used to construct a phylogenetic tree along with the 18S rRNA gene of PLY432. The 18S rRNA gene of six species from *Saccharomyces* was likewise retrieved and used as outgroup. Details of the genes are listed in **Supplementary Table 10**. Sequences were aligned using MUSCLE (v3.8.31) in MPI Bioinformatics Toolkit with non-aligned regions removed ^82–84^. The phylogenetic tree was constructed via IQ-TREE 2 ^85^ under the automatic optimal model selection (Tne+I+G4), calculated with 1,000 bootstrap replications and visualized with TVBOT ^86^.

### Total carbohydrate quantification

The total carbohydrate content in *Glossomastix* cultures was measured continuously during growth using the phenol-sulfuric acid method ^87^. Briefly, 0.2 mL sample, 5% phenol and concentrated sulfuric acid was combined in a 2 mL tube (1:1:5) and gently mixed. After 10 min at room temperature, the samples were placed in a water bath at 30°C for 20 minutes. The content of each tube was cooled to room temperature and transferred to cuvettes, and the absorbance of each sample was then detected at OD_490_ using a BioSpectrometer (Eppendorf AG) or 100 μL of sample in a 96-well plate using the SpectraMax iD3 plate reader. A calibration curve was constructed by analyzing the linear relationship between the concentration and optical density at 490 nm of the 99.99% standard glucose stock in the range of 0.02-0.5 mg/mL.

### Extraction of total polysaccharides from *Glossomastix*

After two months of cultivation, EDTA was added to 1 L *Glossomastix* culture (50 mM final concentration) and autoclaved (121°C, 15 minutes). Whatman^®^ glass microfiber filters (Grade GF/F, 0.70 μm) and Millipore Express PLUS Membranes (0.22 μm) were used to separate polysaccharide fractions from the supernatant sequentially. The filtered culture supernatant was concentrated on an Amicon^®^ stirred cell (Millipore) equipped with a 30 kDa ultrafiltration membrane to collect the HMW (High Molecular Weight) polysaccharide fraction. The concentrate was continuously dialyzed with ultra-pure water (UPW) until the conductivity of filtrate did not change. The conductivity was measured by SevenCompact Duo S213-Meter. The final concentrate was collected and made up to 100 mL with UPW followed by stirring overnight to detach polymers from the membrane. The desalted samples were lyophilized and stored at room temperature until further analysis. Further separation of high purity polysaccharides was carried out using anion exchange chromatography and size exclusion chromatography **(Fig. S5)**.

### Anion exchange chromatography (AEX)

Crude polysaccharide extracts were further concentrated and purified by anion-exchange chromatography on a XK 26/40 column packed with 90 mL ANX FF resin. The packed column attached to an ÄKTA pure system was first equilibrated with Tris-HCl buffer (50 mM, pH 7.5, degassed) at 5 mL/min, followed by sample application. Crude sample in 100 mL Tris-HCl buffer (1 g/L) was filtered (0.22 μm) to remove insoluble materials and then loaded onto the equilibrated resin. Following sample injection, the column was washed with two column volumes of Tris-HCl buffer to remove unconsolidated fractions followed by two column volumes of Tris-HCl buffer containing 0.5 M NaCl. Finally, fraction collection started immediately using two column volumes of Tris-HCl buffer containing 5 M NaCl as elution buffer to wash the column. The eluates were concentrated and desalted via a Amicon^®^ stirred cells with 30 kDa ultrafiltration membrane as described above and lyophilized.

### Size exclusion chromatography (SEC)

Final purification of polysaccharide was performed using size exclusion chromatography on two HiPrep^TM^ 16/60 Sephacryl^TM^ S-400 HR (120 mL per column) connected in series to a Knauer FPLC system (Azura Bio Purification System) equipped with a refractive index detector. Lyophilized sample (100 mg after AEX) was dissolved in 2 mL Tris-HCl buffer, filtered (0.22 μm) and loaded onto the columns that were equilibrated with 300 mL Tris-HCl buffer (50 mM, pH 7.5, degassed) at a rate of 1 mL/min before sample injection. Columns were eluted with 300 mL Tris-HCl buffer and the polysaccharide containing fractions were pooled, desalted and concentrated via Amicon^®^ stirred cells as above.

### Quantification of monosaccharides with HPAEC-PAD

For the quantification of monosaccharides, samples were analyzed on a ICS-5000+ (Dionex) with pulsed amperometric detection (PAD) equipped with a CarboPac PA10 analytical column (2 × 250 mm) and a CarboPac PA10 guard column (2 × 50 mm) ^88^. In brief, 200 µL lyophilized pure polysaccharide samples (1 mg/mL) or 200 μL microalgae culture (with cells) were hydrolyzed with 200 µL 2 M HCl at 100°C for 24 hours in a pre-combusted (450°C, 4 hours) vial. Supernatants from 100 µL bacterial cultures were hydrolyzed with 100 µL 2 M HCl. After complete acid hydrolysis, 100 µL *Glossomastix* culture samples were dried by speed vacuum to remove HCl and then resuspended in 100 µL UPW, followed by a 1:100 (v/v) dilution. The other samples were diluted with UPW at a ratio of 1:200 (v/v) and then centrifuged at 14,800 rpm (Thermo Scientific™ Fresco™ 21 Microcentrifuge) for 10 minutes. Note that after acid hydrolysis, *Glossomastix* culture samples were dried by speed vacuum and then resuspended in 100 µL UPW, followed by a 100-fold dilution.

Subsequently, 100 µL supernatant was analyzed by direct injection onto the HPAEC-PAD system. Monosaccharide standard **(Supplementary Table 11)** mix ranging from 1-10 to 1000 μg/L were used to identify peaks by retention time and to construct standard curves for quantifying the amount of monosaccharide products in the reaction mixture.

### Desulfation of fucan

A complete desulfation of fucan was conducted using a modified version of the solvolytic desulfation protocol outlined by Inoue and coworkers ^89^. First, sodium cations were exchanged with pyridinium ions by dissolving 20 mg of fucan in water and passing the solution through an AG^®^ 50W cation exchange resin (Bio-Rad), pre-equilibrated with pyridine (Sigma-Aldrich). The eluate was neutralized with 0.3 mL pyridine and subjected to lyophilization. Subsequently, the fucan-pyridinium salt was dissolved in 15 mL of DMSO (Sigma-Aldrich), and 75 µL of UPW was introduced. The mixture was incubated at 80 °C for 30 minutes and then subjected to dialysis (8K molecular weight cut-off) against 1 M NaCl and UPW prior to lyophilization.

### NMR characterization of *Glossomastix* fucan

SEC-purified fucan is used only for structural and elemental analysis. Purified fucan and desulfated fucan (10 mg) were first dissolved in 1 mL 99.9% D_2_O (Sigma-Aldrich) and lyophilized in order to reduce the residual water signal. Subsequently, the samples were dissolved in 200 μL D_2_O (D-99.96%; Sigma-Aldrich) and transferred to a 3 mm LabScape Stream NMR tube (Bruker LabScape). For NMR analyses all homo- and heteronuclear experiments were acquired on a Bruker AV-IIIHD 800 MHz spectrometer (Bruker BioSpin AG, Fälladen, Switzerland) equipped with a 5 mm cryogenic CP-TCI z-gradient probe. Chemical shifts were calibrated using the residual water signal for ^1^H (4.75 ppm at 25 °C). The ^13^C chemical shift was internally referenced to DSS using absolute frequency ratio ^90^ (^13^C/^1^H = 0.251449530). For chemical shift assignment the following one- and two-dimensional NMR experiments were recorded at a temperature of 25°C: 1D proton with water suppression (zgesgp), ^1^H-^13^C HSQC with multiplicity editing (hsqcedetgpsisp2.3), ^1^H-^13^C heteronuclear two bond correlation (H2BC) spectroscopy (h2bcetgpl3pr), ^1^H-^13^C heteronuclear multiple bond coherence (HMBC) with suppression of one-bond correlations (hmbcetgpl3nd), ^1^H-^1^H in-phase correlation spectroscopy (IP-COSY) with water suppression with excitation sculpting (ipcosyesgp-tr) ^91^, ^1^H-^1^H total correlation spectroscopy (TOCSY) with 70 ms mixing time and water suppression (clmlevphpr), ^1^H-^13^C HSQC-TOCSY with 70 ms mixing time protons (hsqcdietgpsisp.2), ^1^H-^1^H nuclear Overhauser effect spectroscopy (NOESY) with 80 ms mixing time and water suppression with excitation sculpting and gradients (noesyesgpph). The spectra were recorded, processed, and analyzed using the TopSpin 3.5 or 4.0.1 software (Bruker BioSpin AG).

### Sulfate quantification

The released sulfate from acid-hydrolyzed polysaccharide was measured on a Metrohm 761 compact ion chromatograph equipped with a Metrosep A Supp 5 column and suppressed conductivity detection with 0.5 M H_2_SO_4_. Ions were separated by an isocratic flow of carbonate buffer (3.2 mM Na_2_CO_3_, 1 mM NaHCO_3_) and the duration of each run was 20 minutes, with sulfate eluting at 16 minutes. Chromatograms were analyzed with the instrument’s software MagIC Net version 3.2.

### Elemental analysis

Elemental analysis was performed using an Elementar modern elemental analyzer. Lyophilized sample (0.1-1.0 mg) was transferred to a dry and pre-weighed tin boat and a small amount of tungsten oxide was added. For the calibration curve, sulfanilamide (0.1-1 mg) was prepared in the same way. Before each test, the samples were degassed by compressing tin boats.

### ELISA

Enzyme-linked immunosorbent assay (ELISA) was used to assess the binding of four fucan-specific rat monoclonal antibodies (mAbs), namely BAM1, BAM2, BAM3 and BAM4 ^35^, to *Glossomastix* fucan purified by AEX. The purified fucan was dissolved in water and underwent dilution in PBS to 200 μg/mL followed by five two-fold dilutions in PBS, resulting in a total of six concentrations. Each antibody-fucan combination was tested in triplicate. For the ELISA, 100 μL of the fucan solution were added to wells of a 96-well plate (NUNC MaxiSorpTM, Thermo Fisher Scientific). After overnight incubation at 4℃, wells were washed six times with tap water, and unbound sites were blocked with 200 μL of PBS buffer (137 mM NaCl, 2.7 mM KCl, 10 mM Na_2_HPO_4_, 1.7 mM KH_2_PO_4_, pH 7.4) containing 5% w/v low-fat milk powder (5% MPBS) for 2 hours at room temperature. After washing nine times with tap water, 100 µL of the mAbs diluted 1:10 in 5% MPBS were added to each well and incubated for 1.5 hours at room temperature. After washing nine times with tap water, 100 µL of anti-rat secondary antibody conjugated to horseradish peroxidase (A9037, Sigma-Aldrich) diluted 1:1000 in 5% MPBS were added to each well and incubated for 1.5 hours at room temperature. Wells were washed with tap water nine times. The plate was developed by adding 100 µL ELISA tetramethylbenzidine (TMB) substrate per well. The enzyme reaction was stopped after 5-10Dminutes by addition of 100 µL 1 M HCl to each well. Absorbance at 450 nm (mAb binding intensities) was measured with a SPECTROstar^®^ Nano absorbance plate reader using the MARS software (BMG Labtech). Fucan from *Laminaria* (Glycomix, PSa13) was used as a positive control. Appropriate negative controls were run on every plate.

### Media and monitoring of growth

Unless otherwise stated, bacteria were cultivated in marine minimal Tris-HCl medium ^92^ supplemented with vitamins ^93^, and iron. The base minimal medium (MMT) contained 2.3% (w/v) sea salts (S9883, Sigma-Aldrich), 9 mM NH4Cl, 26 mg/L ammonium ferric citrate (F5879, Sigma-Aldrich), 50 mM Tris-HCl (pH 7.8), and 1 × vitamin mix (1 L 1000 × vitamin mix: 10 mg biotin, 10 mg lipoate, 50 mg Ca-D-pantothenate, 50 mg vitamin B12, 100 mg nicotinate, 100 mg pyridoxamine dihydrochloride, 100 mg thiamine hydrochloride, 40 mg aminobenzoate and 30 mg folate). MMT was further supplemented with 0.03% (w/v) yeast extract (MMT-YE), 0.03% (w/v) casamino acids (MMT-CA), or 50-100 µM KH_2_PO_4_ (MMT-KDP). At a concentration of 0.03% (w/v) BactoTM casamino acids, the medium already contains ∼ 48.9 μM Pi ^94^. All media were sterilized by 0.22 µm filtering. Unless otherwise stated, bacteria were grown at room temperature (400 rpm) in 24-well plates (Sarstedt, 83.3922.500) and growth was monitored as the optical density at 600 nm (OD_600_) using a SpectraMax iD3 plate reader (Molecular Devices).

### Bacteria and growth conditions

‘*Lentimonas*’ sp. CC4 was obtained previously ^5^ and *Wenyingzhuangia fucanilytica* CZ1127^T^ (DSM 100787) was purchased from DSMZ (German Collection of Microorganisms and Cell Cultures). Bacteria were isolated from non-axenic *Glossomastix* by plating 1:10^4^ diluted microalgal culture onto Difco^TM^ Marine Agar 2216. Single colonies were obtained from plates after incubation at 15°C for 12 days. All bacteria were cultured in MMT-CA medium containing 0.05% (w/v) glucose for 5 days, and then inoculated 1:1000 (v/v) into MMT-CA medium containing 0.05% (w/v) *Glossomastix* fucan. OD_600_ values of the cultures were measured on day six and ability to degrade fucan was assessed by C-PAGE as described below.

### Isolation of fucan-degrading bacterium *V*_227

Sediment-associated bacteria were sampled on 27th February 2022 at low tide from the North Beach of Helgoland, Germany, and used to inoculate 13 mL polypropylene tubes (Sarstedt, 62.515.006) containing 3 mL MMT-YE medium supplemented with 0.2% (w/v) fucan (Glycans eluted in AEX using 0.5 M NaCl). Enrichment cultures were incubated at 17°C, 115 rpm for two weeks, and then diluted 1:100 (v/v) into fresh MMT-YE medium with fucan (AEX 5 M NaCl fraction) and cultivated for additional 7 days. Cultures showing an increase in optical density were diluted 1:10^5^ with fresh MMT medium and plated on solid MMT-YE with 0.05% (w/v) fucan and 1% (w/v) agarose. After incubating at room temperature for 5 to 6 days, the putative fucan-degrading isolates appearing as colonies on the plates were re-inoculated into fresh MMT-YE medium to confirm growth and degradation of fucan.

### Detection of fucan degradation

Bacterial degradation of fucan in the culture medium was analyzed qualitatively by the BSA-acetate method ^95,96^ or C-PAGE ^97,98^, and quantitatively by toluidine blue (TB) assay ^99^.

#### BSA-Acetate method

For the BSA-acetate assay, 20 µL culture supernatant was mixed in a 96-well plate with 180 µL acid albumin solution (per liter: 3.26 g of sodium acetate, 4.56 mL of glacial acetic acid, and 1 g of bovine serum albumin, pH adjusted 3.72 to 3.78). Degradation of fucan was assessed by observing the formation of cloudy white precipitates against a black background; the degree of turbidity correlates positively with the concentration of acidic polysaccharides and thereby a clear transparent solution indicated fucan degradation.

#### Carbohydrate polyacrylamide gel electrophoresis (C-PAGE)

Bacterial cultures with fucan were centrifuged at 12,000 rpm (Thermo Scientific™ Fresco™ 21 Microcentrifuge) for 10 min, and 24 µL of supernatant was mixed with 6 µL of 5 × Phenol Red loading dye before loading onto an acrylamide gel (25 % resolving/5 % stacking). Electrophoresis was performed for 30 min at 100 V followed by 1 hour at 200 V in native running buffer (1 L: 3 g Tris 15 g glycine). The gel was stained with 0.005% (w/v) Stains-All in 30% ethanol overnight and de-staining with UPW until the background of the gel was clear.

#### TB method

For the TB assay, 10 µL culture supernatant was mixed with 990 µL TB solution (0.03 mM toluidine blue in 20 mM citrate buffer, pH 3.0, 0.22 µm filtered). Then, 100 μL solution was transferred to a 96-well plate and absorbance measured at 632 nm in a SpectraMax iD3 plate reader (Molecular Devices). Absorbance values were converted to concentration via standard curves constructed on *Glossomastix* fucan (0 to 1 mg/mL) where sulfated fucan concentration is inverse proportional optical density 632 nm **(Supplementary table 12)**.

### Growth physiology of *V*_227

#### Carbohydrate assimilation experiments

Growth assays with mono- and polysaccharides were performed in multi-well culture plates (0.5-1 mL medium) or 13 mL culture tubes (4 mL medium) depending on the availability of carbohydrate substrate and volume needed for glycan detection. For growth assays with *Glossomastix* fucan, a 4 day old preculture of *V*_227 grown in MMT-YE was inoculated 1:100 (v/v) into 4 mL fresh MMT-YE medium (20°C, 110 rpm). Negative control means that no carbohydrates were added. This growth experiment was performed in 13 mL polypropylene tubes (Sarstedt, 62.515.006). Sampling was performed at intervals of 6 to 14 hours and the OD_600_ was measured in 10 mm cuvettes using a BioSpectrometer (Eppendorf AG). Supernatants were centrifuged at 12,000 rpm (Thermo Scientific™ Fresco™ 21 Microcentrifuge) for 10 minutes and collected for fucan detection (all time points) and monosaccharides detection (some time points) after acid hydrolysis via HPAEC-PAD. This growth experiment was performed in test tubes. In subsequent growth experiments, pre-cultures were inoculated into fresh medium at a ratio of 1:1000. Monosaccharides **(Supplementary Table 11)** used in the growth assay at a final concentration of 0.05% (w/v). For the fucose, rhamnose, galacturonate and glucuronate, *V*_227 was incubated for three weeks (MMT-CA, 24-well plate or test tubes, 20°C, 110 rpm). For glucose and galactose, *V*_227 was incubated for one week (MMT-KDP, 24-well plate, RT, 400 rpm).

Growth experiments with *Fucus vesiculosus* fucan and *Glossomastix* fucan as carbohydrates were carried out in MMT-CA medium at a final concentration of 0.05% (w/v) and OD_600_ was measured at 6 to 24 hours intervals (24-well plate, RT, 400 rpm). Negative control means that no carbohydrates were added. We simultaneously tested the growth of *V*_227 with the addition of different polysaccharides in MMT-KDP medium (100 µM KH_2_PO_4_) **(Supplementary Table 11)**, and the cultivation was performed in 48-well plates (Sarstedt, 83.3923.500) (RT, 400 rpm). OD_600_ was measured via BioSpectrometer (Eppendorf AG) after one week incubation.

#### Phosphate-limitation experiments

*V*_227 was cultivated in MMT-KDP medium (1–50 µM KH_2_PO_4_ final concentration, for pectin is 1-30 μM) with 0.05% (w/v) of polysaccharide as the sole carbon source: *Glossomastix* fucan (24-well plate, RT, 400 rpm), *Eisenia bicyclis* laminarin (24-well plate, RT, 400 rpm), Corncob xylan (24-well plate, RT, 450 rpm) and *Sugar beet* pectin (24-well plate, RT, 450 rpm) **(Supplementary Table 11)**. Growth was monitored during 144 hours of incubation after which, the cultures were centrifuged at 12,000 rpm (Thermo Scientific™ Fresco™ 21 Microcentrifuge) for 10 min. Supernatants were subjected to monosaccharide composition analysis via HPAEC-PAD and total carbohydrate quantification as described above. The converted glycan carbon is the sum of the initial molar concentrations of all monosaccharides minus the sum of the molar concentrations of all monosaccharides remaining in the supernatant at 144 hours, and then the value obtained is multiplied by 6 for these hexoses having six carbon atoms.

### Calibration of optical density against cell counts

*V*_227 was grown in MMT-CA medium with 0.05% (w/v) fucan. When *V*_227 was in the logarithmic growth phase (OD_600_ = 0.262), cultures were diluted into groups with different cell densities. The OD_600_ of the diluted cultures was measured in 24-well plates using a SpectraMax iD3 plate reader. Each serial dilution was then further diluted 10,000 times and 100 μL was spread on MMT-CA plates (0.05% fucan and 1% agar) and incubated for 9 days. Linear correlation analysis was performed between the number of colonies counted and the OD_600_ value **(Supplementary Table 13)**.

### Super-resolution microscopy of selfish fucan uptake

Fucan was fluorescently labelled with fluoresceinamine (Sigma, 3326-34-9) as previously described ^100^. Precultures of *V_*227 were diluted 1:1000 in 1 mL MMT-KDP medium (50 μM KH_2_PO4) containing 0.4% (w/v) fluorescently labelled fucan and triplicate cultures were grown in a 24-well plate (RT, 400 rpm). At several time points over 7 days, 100 μL culture was fixed with 2% (v/v) formaldehyde and diluted 1:10 in 1 × PBS buffer. The fixed cells were collected on polycarbonate filters (0.2 μm, Ø47 mm) before staining with 4′,6-diamidino-2-phenylindole (DAPI) and then mounted onto glass slides using a Citifluor/VectaShield (4:1) solution. Likewise, 100 μL *V*_227 pre-culture and 100 μL MMT-KDP medium was fixed, DAPI-stained and applied to glass slides as controls. Stained *V*_227 cells (day 3) were visualized by epifluorescence microscopy on an AxioImager.Z2 microscopic stand (Carl Zeiss), equipped with automated imaging and light-emitting diodes ^101,102^. Images were acquired at 63 × magnification. For super-resolution microscopy the cells were visualized on a Zeiss ELYRA PS.1 (Carl Zeiss) using 488, and 405 nm lasers and BP 502-538, and BP 420-480 + LP 750 optical filters. Zstack images were taken with a Plan-Apochromat × 63/1.4 oil objective and processed with the software ZEN2011 (Carl Zeiss, Germany).

### Genome sequencing and assembly

Genomic DNA was extracted with the Gentra Puregene Yeast/Bact. Kit (Qiagen) using 5 mL culture of *V_*227 which was grown in MMT-YE medium (20°C, 110 rpm) for 4 days and the purified genomic DNA was then sequenced on a PacBio Sequel II platform at the Max Planck Genome Centre Cologne. An assembly of *V_*227 genome was carried out using HiCanu v2.2 ^103^ based on PacBio hifi reads and assembly quality evaluated by checkM v1.2.2 ^104^.

### Relative abundance in TARA and Helgoland spring bloom water

We performed read mapping against the TARA read dataset using Bowtie2 v2.3.5.1 ^105^. Samtools v1.7 ^106^ was used to convert the SAM files to BAM, which were then sorted. The trimmed_mean value across all reads were obtained from the sorted BAM files via CoverM v0.6.1 (https://github.com/wwood/CoverM). The trimmed mean was normalized using the genome equivalence of the *RpoB* gene as was calculated from the TARA dataset. This was then repeated using the Helgoland spring bloom read dataset (Mid-March ∼ Mid-May of 2010, 2011, 2012, 2016, 2018, and 2020) ^107,108^, except that the trimmed mean was then normalized using the genome equivalence of the *RpoB* gene as was calculated from the Helgoland dataset. The genome equivalence of the *RpoB* gene was obtained by mapping all reads in the TARA dataset against a collection of reference *RpoB* sequences ^46,109^ using the same tools detailed above. The genome equivalence for each sample was calculated as follows: (total number of hits to the *RpoB* references × average read length) ÷ the average length of the *RpoB* references. This analysis was then repeated using the Helgoland dataset.

### Phylogenetic tree of novel fucan-degrading *Verrucomicrobiaceae* sp. isolate

Genomes and MAGs belonging to the *Akkermansiaceae* family were obtained from GTDB v214.1 ^110^. Genomes used were selected based on their quality (completion – 5 x contamination >= 50) as determined using checkM v1.2.2 ^104^ and then dereplicated using a 99% ANI threshold for the secondary clustering step in dRep v3.4.3 ^111^. A total of 246 selected genomes **(Supplementary Table 7)** were further used for phylogenetic reconstruction based on 120 conserved genes determined using GTDB-tk v2.3.2 ^112^ and the v214.1 GTDB release. A maximum likelihood tree was determined using IQ-TREE v2.2.2.6 (-m MFP) ^85^ and visualized using the interactive Tree of Life (iTol) ^113^.

### Genome annotation

Preliminary annotation of the *V*_227 genome was performed with Prokka (v1.14.5) ^114^. The proteins domain families were annotated using Pfam-A HMMs ^115^ via HMMER (v3.3.2, http://hmmer.org) with a “cut_tc” thresholding, and the hit with the highest score value for each sequence was extracted. CAZymes were predicted using a combination of HMMER (E-value < 1e−15, query coverage > 35%) against the dbCAN v11 (https://bcb.unl.edu/dbCAN2/download/Databases/V11/) HMM database and Diamond blastp ^116^ (v2.0.14.152, E-value < 1e-20, identity > 40% and query coverage > 50%) against the 2022 CAZy database ^59^. Only results with consistent dbCAN and CAZy annotations were considered reliable. Sulfatases were annotated based on Diamond blastp (v2.0.14.152, E-value < 1e-20, identity > 40% and query coverage > 50%) searches against the SulfAtlas database v1.3 ^117^. Transporters were annotated by Transporter Automatic Annotation Pipeline (TransAAP) ^118^. Alpha-L-fucosidases were annotated using a reference HMM models (PF01120) via HMMER. The MetaCyc database^119–121^ and BlastKOALA ^122^ were used for metabolic pathway reconstruction and confirmed by InterPro ^123^ or CDD ^124,125^. Prodigal (v.2.6.3) ^126^ was used to obtain amino acid sequences of proteins encoded in 245 bacterial genomes selected from *Akkermansiaceae* family. Proteins in the Phosphate transport high affinity-system (PstA, PstB, PstC and PstS) were identified in the 245 genomes via HMMER using the individual HMM modules: PstA (TIGR00974.1, E-value < 1e-15, query coverage > 50%), PstB (TIGR00972.1, E-value < 1e-50, query coverage > 50%), PstC (TIGR02138.1, E-value < 1e-15, query coverage > 50%) and PstS (TIGR00975.1, E-value < 1e-05, query coverage > 35%). A bacterium was considered to encode Pst when at least three homologs were identified in its genome.

### Data visualization and statistics

Most data visualizations and statistical analyses were generated using Python with Matplotlib ^127^, Pandas ^128^, Numpy ^129^, Scipy ^130^ and Statsmodels ^131^ packages. The growth rate was obtained using R with Growthcurver ^132^ package. Fig. 3f, extended data fig. 3a, and extended data fig. 4d were created with BioRender.com. Extended data fig. 4a were created with Google Maps.

## Data availability

All relevant data supporting the results of this study are available in the paper and its supplementary information. The 18S rRNA gene sequence of *Glossomastix* sp. PLY432 has been deposited in NCBI under accession number PP265255. The genomic data of *V*_227 has been deposited in NCBI under accession number PRJNA1070871. The raw data of HPAEC-PAD plus the resulting data tables used for calculation are currently being deposited on the Pangaea data repository https://www.pangaea.de/. A digital object identifier for these data will follow. Please note the tables have been also submitted as supplementary files.

## Acknowledgments

We thank Jaagni Parnami and Kirsten Imhoff for sulphate quantification. Alek Bolte, Gabriele Klockgether and Katharina Föll for HPAEC-PAD measurements and elemental measurements. Tina Trautmann for help in pre-treatment of microalgal cultures and *Glossomastix* genomic DNA extraction. Maren Oftebro for desulfating the glycan prior to NMR characterization. Bruno Huettel from Max Planck Genome Centre Cologne for genome sequencing. We thank Ian Probert from Roscoff Culture Collection for providing *Glossomastix* sp. PLY432. Tao Song and Guoyin Huang for advice on experimental design. We are grateful for institutional support from the University of Bremen, MARUM, MPIMM.

## Author Contributions

Y.X., D.Q. and J.H.H. conceived and planned the study. Y.X., Yi.C. and J.H.H. initiated the research. Y.X. and M.S.J. isolated bacteria. Y.X., Yu.C. and B.G. performed growth experiments. Y.X. and S.V.M. performed ELISA. L.J.K. and F.L.A. obtained and interpreted NMR spectroscopy data. I.W., L.H.O. and Y.X. did bioinformatic analyses. M.F. and G.R. did fluorescence labeling and microscopy. Y.X., H.Y. and J.H.H. analyzed the data and designed illustrations. Y.X., H.Y., M.S.J. and J.H.H. wrote the manuscript with input from all authors.

## Extended Data figures

**Extended Data Fig. 1:**
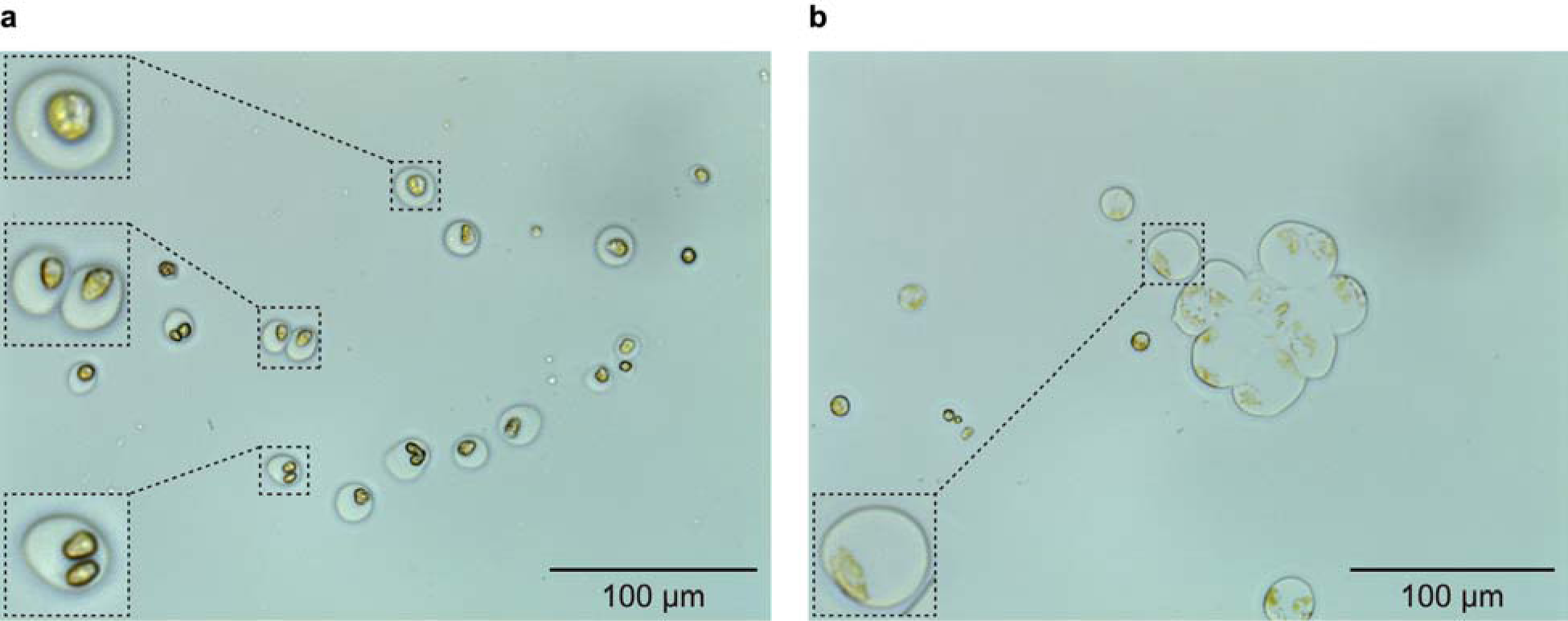
*Glossomastix* cells were coated with mucus. **a,** Microscopic imaging for less viscous *Glossomastix* culture. Some algal cells have finished their bipartition period and are detaching from each other. **b,** Microscopic imaging for highly viscous *Glossomastix* culture. Cell fragments were enveloped in mucus and formed aggregates.

**Extended Data Fig. 2:**
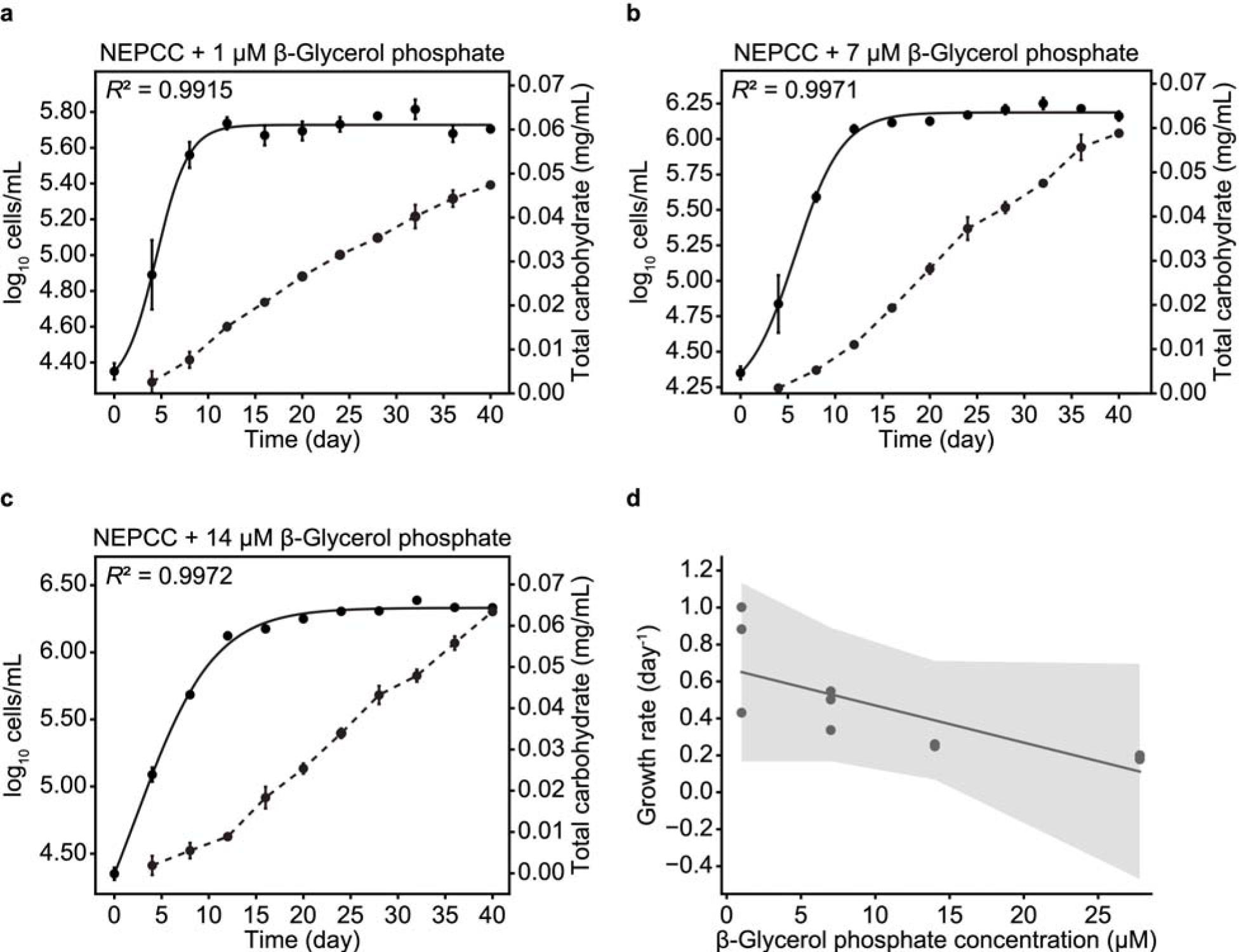
Phosphate concentration shows no significant effect on the growth of *Glossomastix*. **a–c,** Dynamic monitoring of total carbohydrate content and cell density in *Glossomastix* cultures with the addition of different concentrations of β-Glycerol phosphate as a phosphorus source. **d,** Correlation analysis between the growth rate of *Glossomastix* and phosphate concentration (*r* = −0.89, *P* > 0.05). The growth rate for 27.8 µM β-Glycerol phosphate is generated from Fig. 2a. The experiment was performed in independent triplicate (n = 3), error bars are the standard deviation of the mean. The fitting was performed using the mean of triplicate.

**Extended Data Fig. 3:**
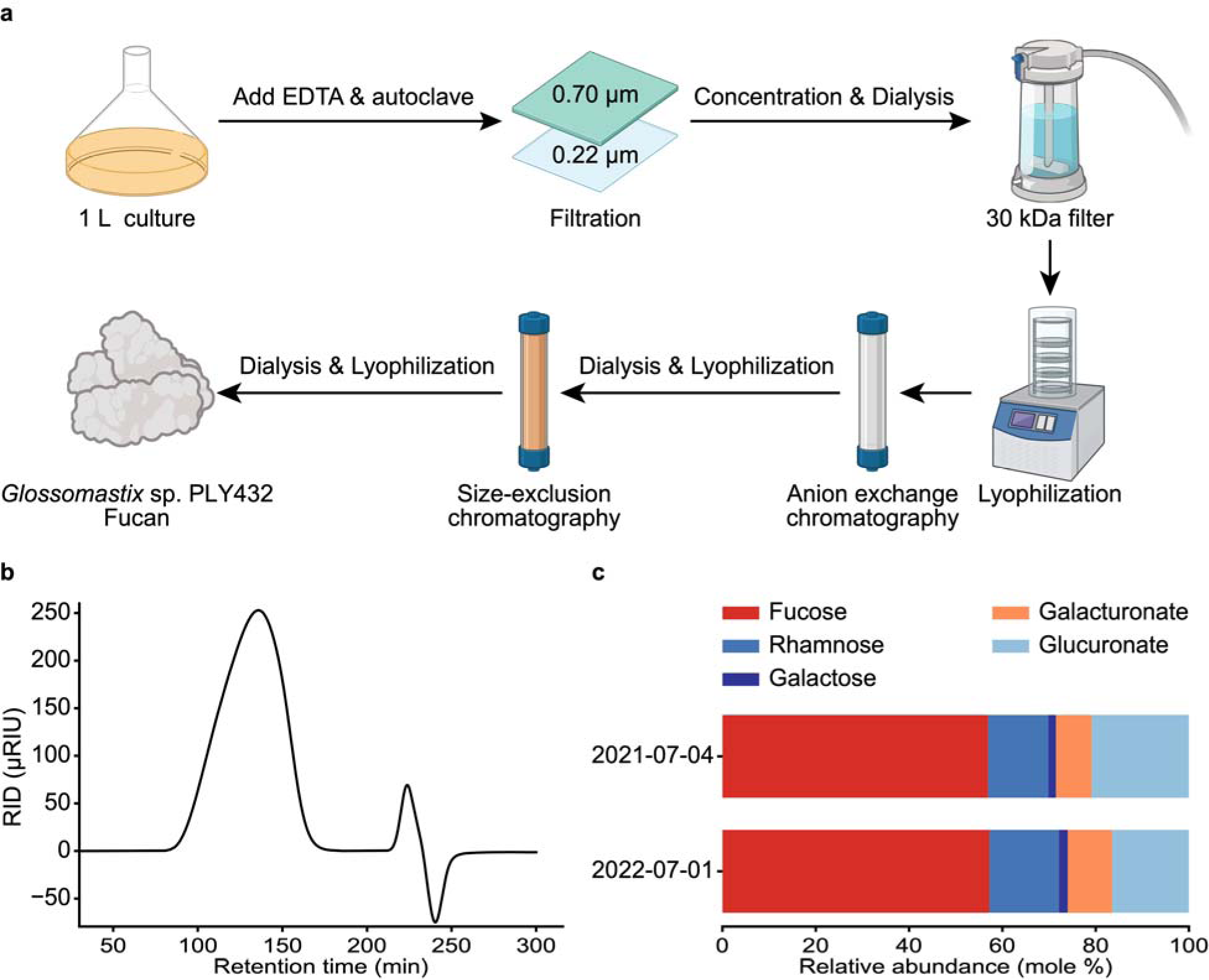
Purification of *Glossomastix* fucan. **a,** Schematic representation of the *Glossomastix* fucan purification workflow. The panel was created with BioRender.com. **b,** Size exclusion chromatography for AEX-purified fucan. **c,** Monosaccharide composition analysis of different, one year apart batches of AEX-purified fucan. The *Glossomastix* cells were continuously cultivated.

**Extended Data Fig. 4:**
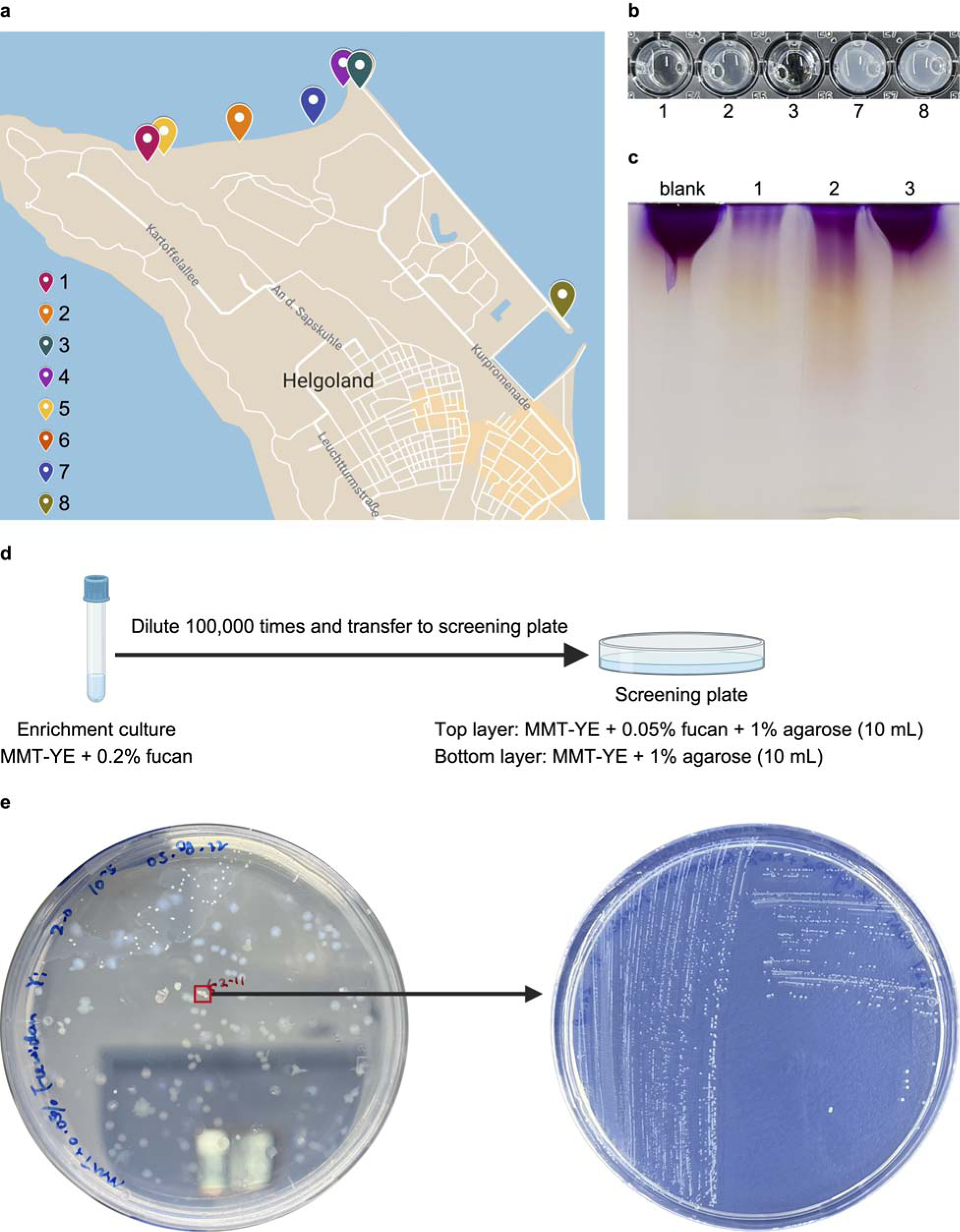
Process of isolating fucan degraders *V_*227. **a,** Sampling location. **b,** Detection of fucan degradation via BSA acetate method. As the clarity of the solution increases, there is a proportional escalation in the degradation of fucan. **c,** Carbohydrate polyacrylamide gel electrophoresis (C-PAGE) for the consumption of polysaccharide. Note that the enrichment 3 in the BSA assay shows degradation of fucan, but after transferring into fresh medium the degradation ability was lost as indicated by the high molecular weight band of fucan on the gel. **d-e,** Isolation steps and the physiological morphology of bacteria on MMT-YE + 0.05% fucan agarose plates.

**Extended Data Fig. 5:**
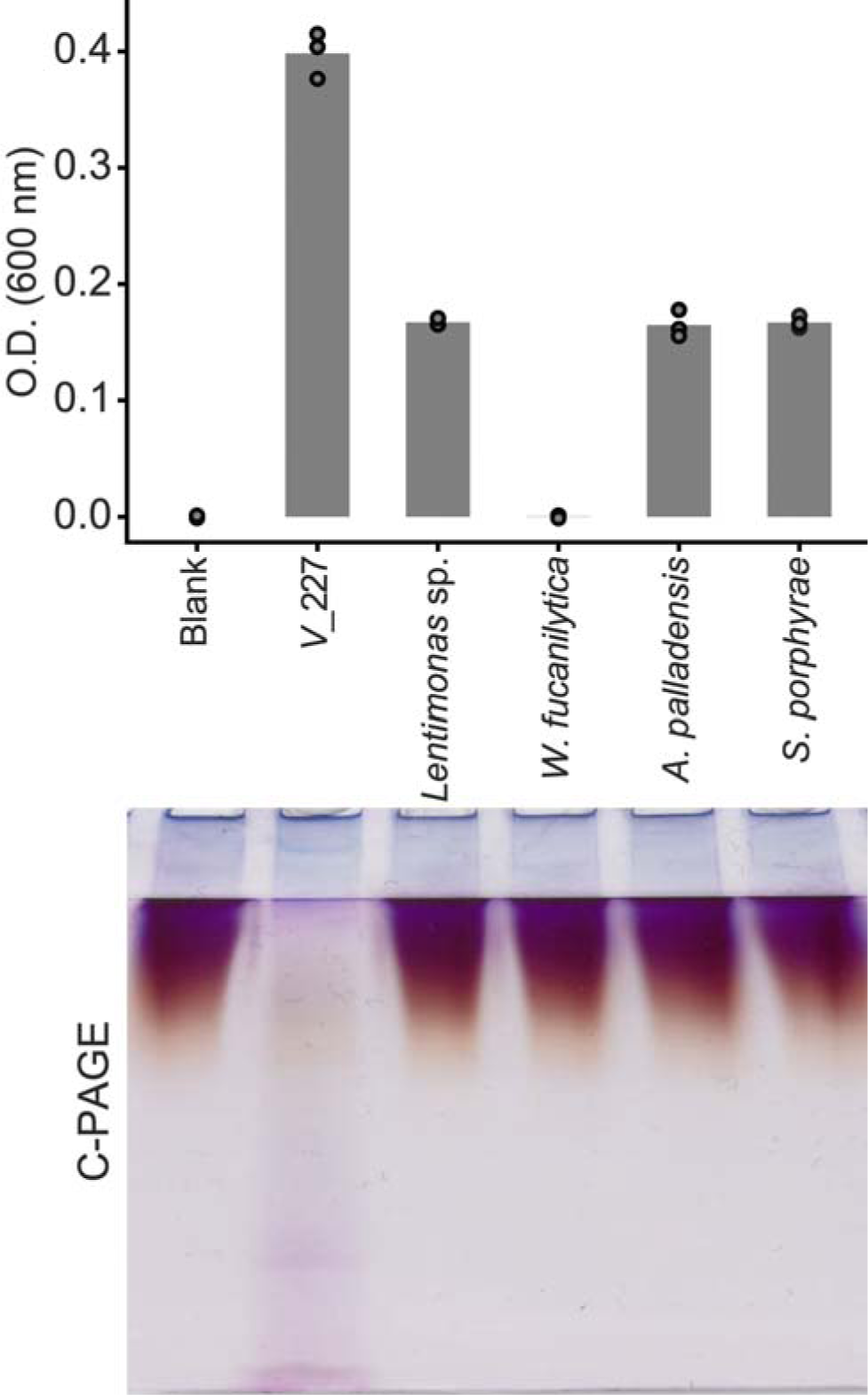
*Verrucomicrobiaceae* bacterium *V_*227 is a specific degrader of *Glossomastix* fucan. Top of the panel, The optical density of the bacteria at 600 nm after 6 days of growth. The bacteria were inoculated in 1 mL MMT-CA medium with 0.5 mg/mL *Glossomastix* fucan at a ratio of 1:1000. *V_*227 in this study is the strain isolated from marine environment. *Lentimonas* sp. CC4 and *Wenyingzhuangia fucanilytica* CZ1127^T^ are well-known degraders of the macroalgae fucan. *Arenibacter palladensis* and *Sulfitobacter porphyrae* are isolated from *Glossomastix* culture. The experiment was performed in independent triplicate (n = 3), error bars are the standard deviation of the mean. Bottom of the panel, carbohydrate polyacrylamide gel electrophoresis (C-PAGE) for the consumption of polysaccharide in the culture supernatant at day 6. One set of samples from three independent replicates is shown.

**Extended Data Fig. 6:**
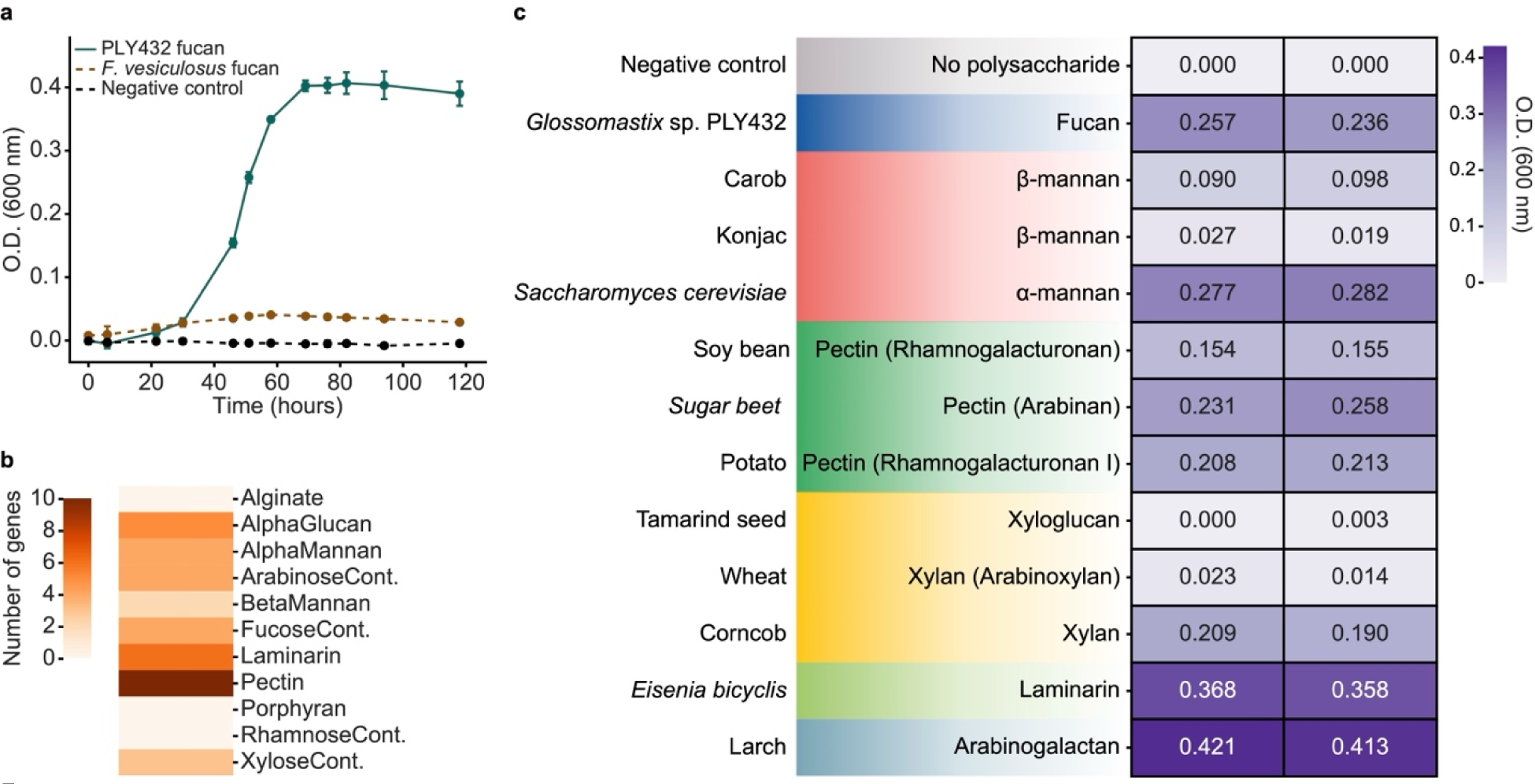
Consumption of different polysaccharides by *V_*227. **a,** Growth of *V_*227 in MMT-CA medium with microalgae and macroalgae fucan. All groups use 0.03% (w/v) Bacto^TM^ casamino acid as the phosphate source, so the medium already contains _∼_ 48.9 μM phosphate. The growth experiment was performed in independent triplicate (n = 3), and error bars are the standard deviation of the mean. **b,** The number of genes targeting different polysaccharides based on CAZy annotations. **c,** Growth of *V_*227 in MMT-KDP medium with different polysaccharides. The experiment was performed in independent two replicates (n = 2).

**Extended Data Fig. 7:**
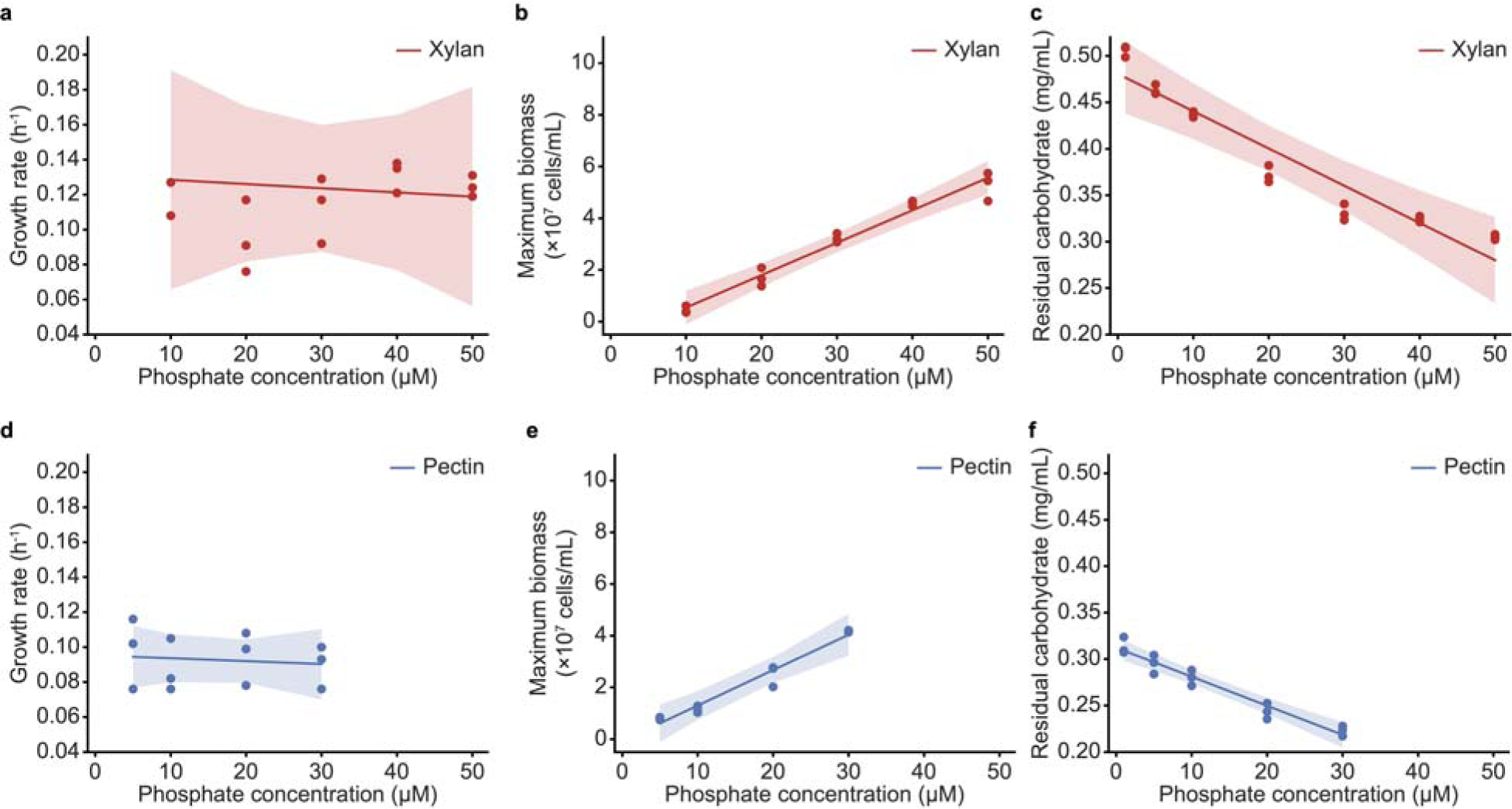
Phosphate limits the utilization of complex carbohydrates by *V_*227. **a,** Correlation analysis between growth rate and phosphate concentration on xylan (*r* = −0.17, *P* > 0.05). **b,** Correlation analysis between maximum biomass and phosphate concentration on xylan (*r* = 0.99, *P* < 0.001) **c,** Correlation analysis between residual carbohydrate in the culture supernatant and phosphate concentration on xylan (slope = −0.004, *r* = −0.96, *P* < 0.001). **d,** Correlation analysis between growth rate and phosphate concentration on pectin (*r* = −0.39, *P* > 0.05). **e,** Correlation analysis between maximum biomass and phosphate concentration on pectin (*r* = 0.99, *P* < 0.01) **f,** Correlation analysis between residual carbohydrate in the culture supernatant and phosphate concentration on pectin (slope = −0.003, *r* = −0.99, *P* < 0.001). All the fitting was performed using the mean of independent triplicate (n = 3).

**Extended Data Fig. 8:**
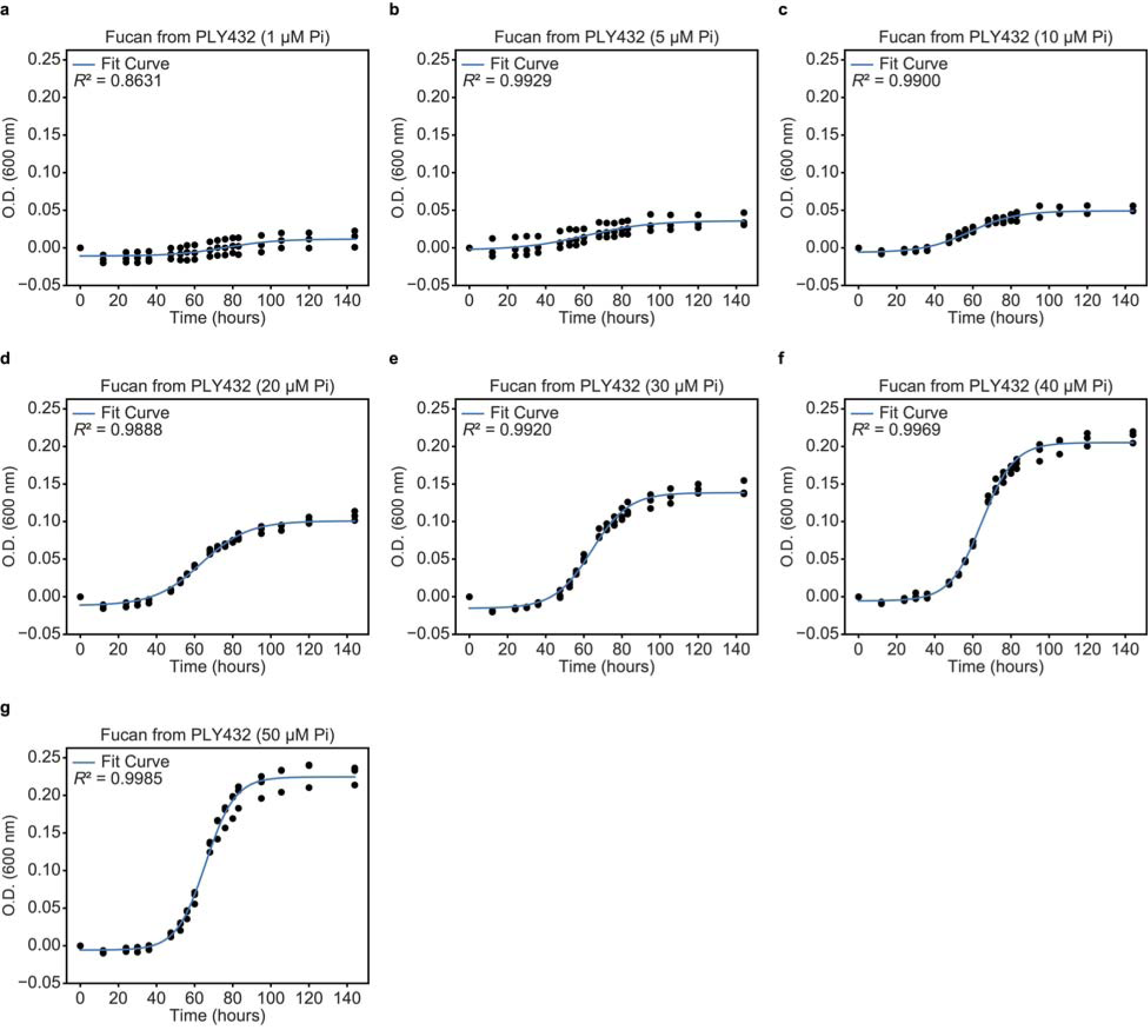
Growth of *V_*227 in MMT-KDP medium containing *Glossomastix* fucan as the carbon source with varying concentrations of phosphate. The experiment was performed in independent triplicate (n = 3). The fitting was performed using the mean of triplicate.

**Extended Data Fig. 9:**
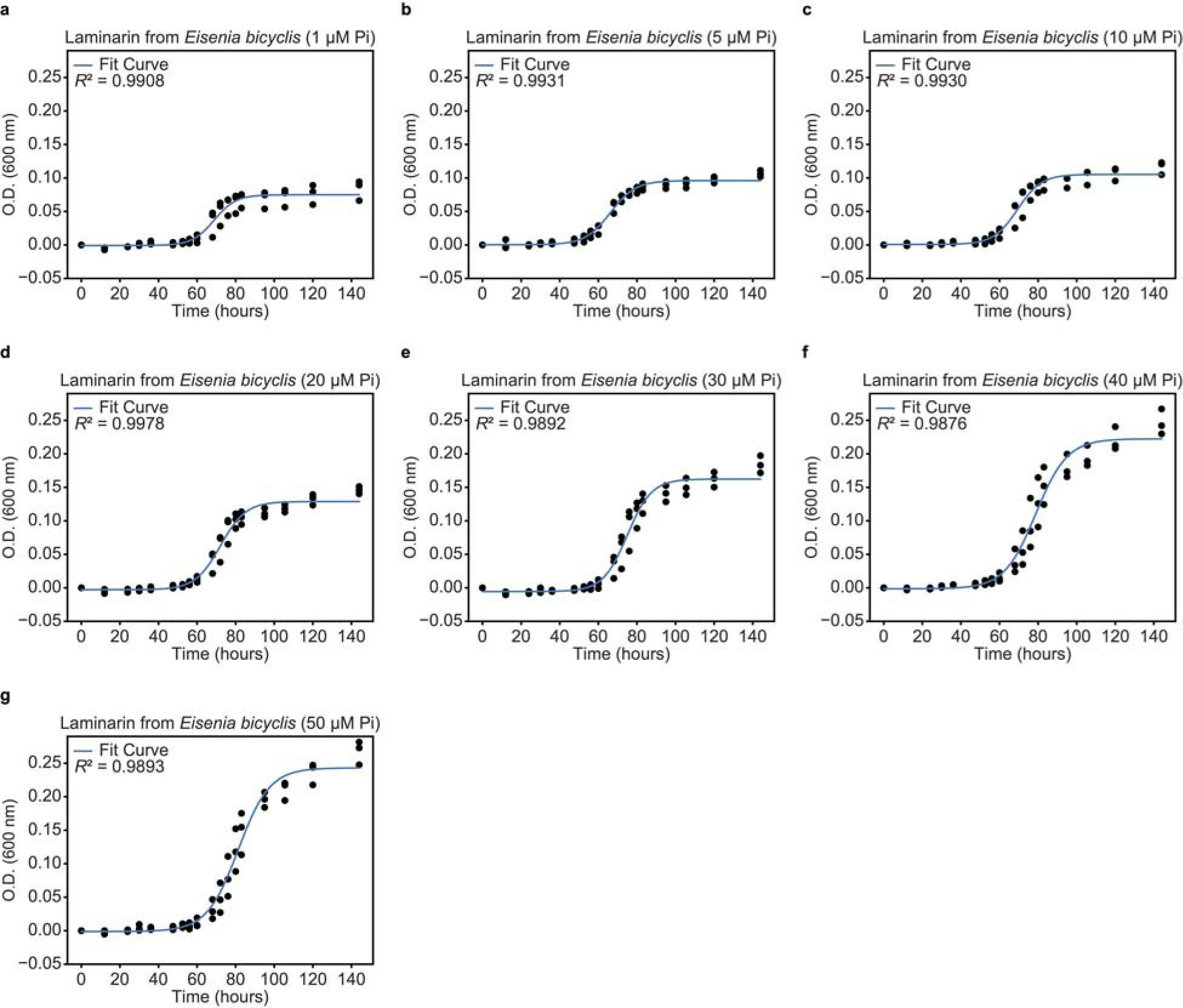
Growth of *V_*227 in MMT-KDP medium containing *Eisenia bicyclis* laminarin as the carbon source with varying concentrations of phosphate. The experiment was performed in independent triplicate (n = 3). The fitting was performed using the mean of triplicate.

**Extended Data Fig. 10:**
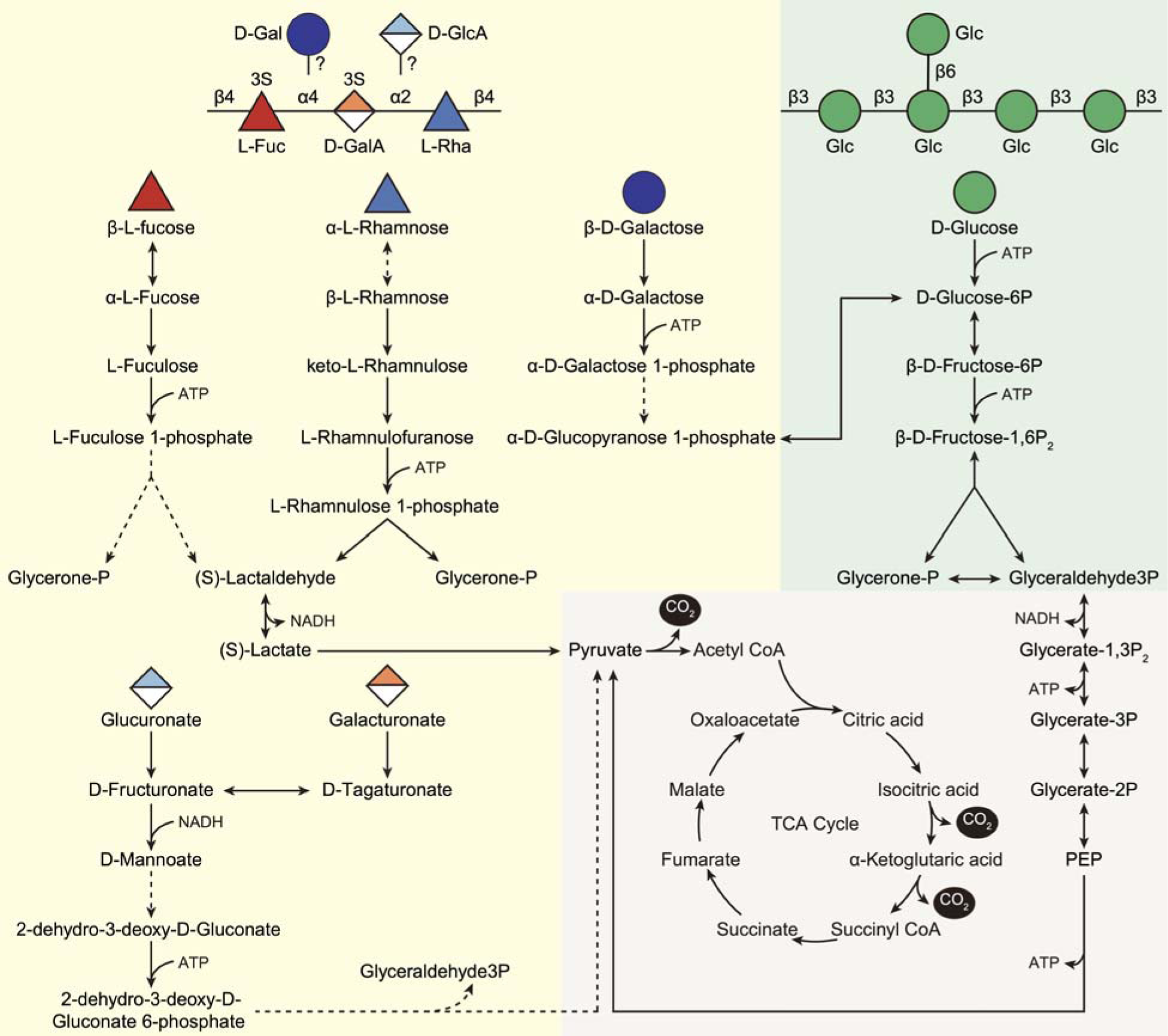
Reconstructed metabolic pathways for six monosaccharides in *V*_227. Solid lines represent homologs found in the genome or this reaction is spontaneous, dashed lines represent homologs not found. The structure of *Eisenia bicyclis* Laminarin and the partial structure of *Glossomastix* fucan are on the top.

## Supplementary figures

**Supplementary Fig. 1:**
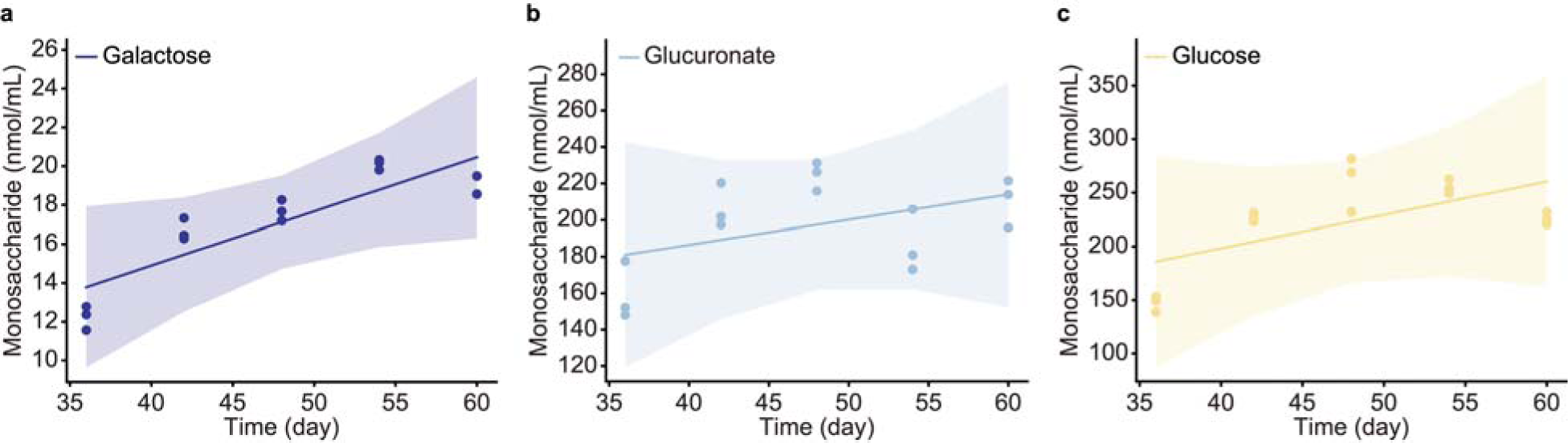
Monosaccharides that did not show significant linear changes in *Glossomastix* cultures from day 36 to day 60. **a–c**, Correlation analysis for galactose (*r* = 0.87, *P* > 0.05), glucuronate (*r* = 0.52, *P* > 0.05) and glucose (*r* = 0.65, *P* > 0.05) in PLY432 cultures along with incubation time. The experiment was performed in independent triplicate (n = 3), the fitting was performed using the mean of triplicate.

**Supplementary Fig. 2:**
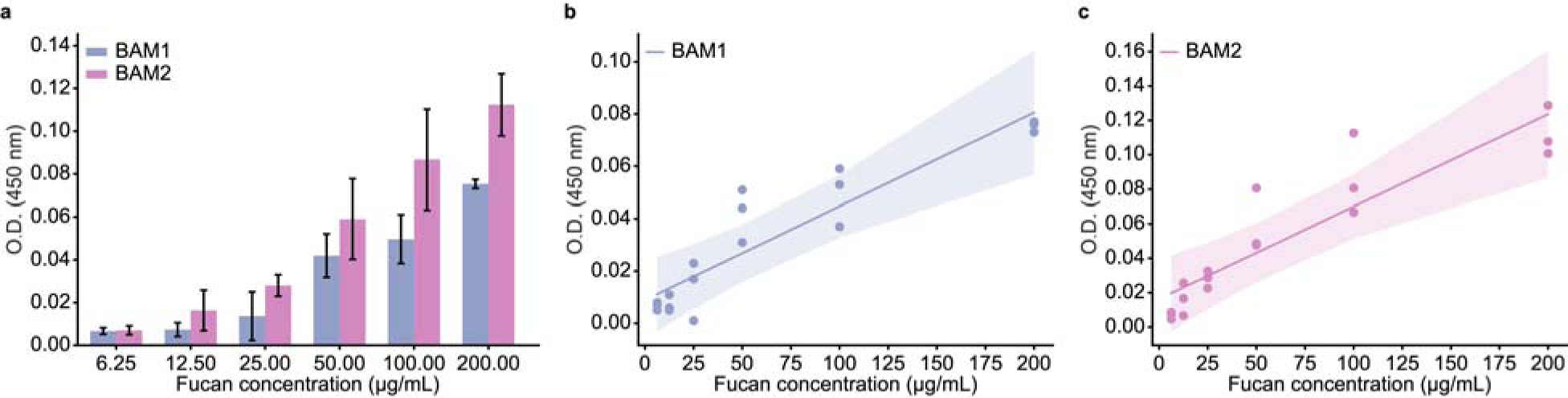
Enzyme-linked immunosorbent assay (ELISA) of *Glossomastix* fucan. **a,** Binding of monoclonal antibodies BAM1 and BAM2 at different concentrations of *Glossomastix* fucan was evaluated with ELISA and binding intensity was read at an OD of 450 nm. **b-c,** Correlation analysis for BAM1 (*r* = 0.95, *P* < 0.01) and BAM2 (*r* = 0.95, *P* < 0.01) with different concentration of fucan. The experiment was performed in triplicate (n = 3), error bars are the standard deviation of the mean.

**Supplementary Fig. 3:**
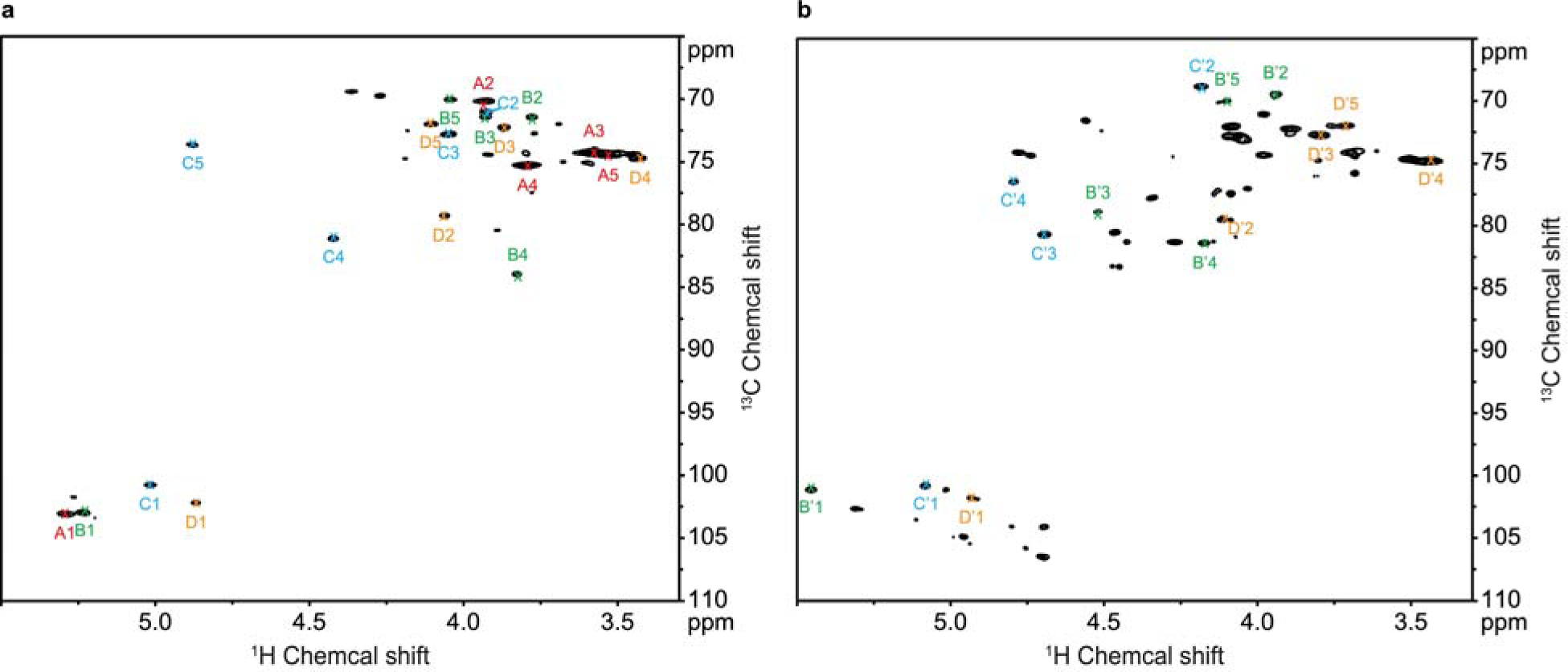
HSQC spectrum for *Glossomastix* fucan. **a,** ^1^H-^13^C HSQC spectrum for desulfated fucan from *Glossomastix*. Correlations are annotated with letter that refers to spin system and number that indicates the carbon number for position within residue. A: α-_D_-GlcA*p* (red); B: α-_L_-Fuc*p* (green); C: α-_D_-GalA*p* (blue) and D: β-_L_-Rha*p* (orange). **b,** ^1^H-^13^C HSQC spectrum for purified fucan from *Glossomastix*. Correlations are annotated with letter that refers to spin system and number that indicates the carbon number for position within residue. B’: α-_L_-Fuc*p* (green); C’: α-_D_-GalA*p* (blue) and D’: β-_L_-Rha*p* (orange). The sample was dissolved in D_2_O (200 µL, 99.96% D), spectrum recorded at 25°C and 800 MHz. ^1^H chemical shift internally referenced to the residual water signal (4.75 ppm) and ^13^C chemical shift referenced indirectly to DSS based on ^1^H/^13^C frequency ratio = 0.251449530.

